# The asthma gut microbiota influences lung inflammation in gnotobiotic mice

**DOI:** 10.1101/2022.08.09.502549

**Authors:** Naomi G. Wilson, Ariel Hernandez-Leyva, Anne L. Rosen, Natalia Jaeger, Ryan T. McDonough, Jesus Santiago-Borges, Michael A. Lint, Thomas R. Rosen, Christopher P. Tomera, Leonard B. Bacharier, S. Joshua Swamidass, Andrew L. Kau

## Abstract

The composition of the gut microbiota in early childhood is linked to asthma risk but the role of the gut microbiota in older patients with established asthma is less clear. Here, we used a cohort of 38 school-aged children (19 with asthma) and 57 adults (17 with asthma) to develop a model that aids in the design of mechanistic experiments in gnotobiotic mice. These experiments show that enterotoxigenic Bacteroides fragilis (ETBF) is associated with increased gut permeability, oxidative stress, and markers of Th17-mediated inflammation in the lungs of mice following ovalbumin sensitization and challenge (OSC). Further, ETBF is enriched in a human population with asthma compared to healthy controls. Our results provide evidence that ETBF has the potential to alter the phenotype of airway inflammation in a subset of patients with asthma outside of early childhood which suggests that therapies targeting the gut microbiota may be helpful tools for asthma control.

## Introduction

Asthma is a common respiratory disease characterized by airway inflammation triggered by an allergic response to environmental antigens. While asthma is predominately associated with T helper (Th)-2 inflammation associated with high levels of cytokines IL-4, IL-5, and IL-13, detailed characterization from clinical studies has revealed substantial heterogeneity in immunopathology of this disease (1). For example, signs of Th17-associated inflammation and oxidative stress are associated with corticosteroid resistance and increased frequency of exacerbations (2) (2, 3) (4). Although disease heterogeneity plays an important role in the prognosis and treatment of asthma, the factors that drive the phenotype of asthma are not well understood.

A potential source of disease variability in the lung lies in the diverse immunologic and metabolic activities of the gut microbiota. Gut microbes have been implicated in the pathology of a range of lung diseases including chronic obstructive pulmonary disease (5), fungal (6) and bacterial pneumonia (7, 8), and, notably, asthma (9). One mechanism by which this phenomenon, termed the “gut-lung” axis (10), affects asthma is through the synthesis of bioactive metabolites. For example, microbe-derived molecules like short-chain fatty acids (9, 11) and 12,13-diHOME (12, 13) are thought to permanently alter the immune system during infancy, a key stage of immune and microbiota maturation, and lead to lifelong susceptibility to allergy and asthma. The role of the gut microbiota during a “critical window” (14) of immune and microbiota co-development (15) is supported by metagenomic surveys that show that the microbiome composition during infancy, but not later in childhood, is associated with the development of asthma later in life (9, 16–18). Additionally, gnotobiotic animal studies have shown that asthma-associated microbes from early childhood can modulate susceptibility to experimental models of allergic airway inflammation (AAI) (9). However, even among older children and adults that are beyond the critical window, individuals with asthma harbor alterations in microbial diversity and composition (19–22) compared to healthy controls.

Few studies have been devoted to identifying and defining the effects of the gut microbiota on lung inflammation in older individuals with asthma. One of these studies has demonstrated that the gut commensal, *Akkermansia muciniphila,* which is reduced in individuals with obesity-associated asthma, can mitigate AAI in mice (22). In contrast, a study of AAI severity in gnotobiotic mice colonized with gut microbiota from school-aged children did not find a difference in markers of allergy between mice colonized with microbiota from healthy children and children with asthma (23). Given these variable results, there remains a need to assess the effect of the gut microbiota on established asthma and to find testable mechanisms for human clinical studies.

This study addresses this need by defining gut microbial compositional differences between individuals with and without asthma and tests the effect of these gut microbiota on lung inflammation in a gnotobiotic mouse model. We recruited a cross-sectional cohort of 38 school-aged (6-10 years) and 57 adult (18-40 years) subjects with or without asthma to identify gut microbes that could influence asthma beyond the critical window. We modeled the interpersonal differences in microbiome composition and selected representative donor fecal samples for further characterization in a gnotobiotic mouse model of asthma. We induced AAI in these gnotobiotic mice to test the effects of gut microbes from donors with or without asthma on inflammation in the lungs. Our results show that while typical markers of allergy were not affected by the gut microbiota (23), asthma-associated microbiomes were more likely to harbor enterotoxigenic *Bacteroides fragilis* (ETBF) which was associated with gut barrier dysfunction, as well as oxidative stress and Th17 responses in the lungs of gnotobiotic mice.

## Results

To investigate if asthma is associated with distinct gut microbial signatures outside of early childhood, we recruited 17 adults and 19 school-aged children with physician-diagnosed, moderate-to-severe atopic asthma along with 40 adult and 19 school-aged healthy controls into the previously described Microbiome and Asthma Research Study (24) (MARS; see demographic summary in Supplemental Table 1; see also Methods). We performed V4-16S rRNA amplicon sequencing of participant stool samples obtained at the patient’s baseline and identified amplicon sequence variants (ASVs) using DADA2 (25) (Supplemental Figure 1A). The alpha diversity of samples from patients with asthma was greater than in healthy controls even while accounting for differences between ages (Figure 1A). To determine how demographic features affected community composition, we performed a PERMANOVA on Bray-Curtis dissimilarity distances between fecal microbiomes (Figure 1B). We found that asthma status, age (26), and race (27, 28), but not adiposity (29), sex (30), smoking history (31), or antibiotic exposure (32) within the past year significantly contributed to the variation in subject gut microbiota composition (Figure 1C), even when accounting for sequencing batch effect and other covariates (Supplemental Table 2). Although age is an important factor in determining asthma phenotype, we did not find the interaction of age and asthma to be significant. While our sample size does not permit us to exclude a weak effect, this suggests that differences in microbiome composition due to asthma are not dependent on age in our cohort. Guided by the PERMANOVA results, we performed differential abundance analysis and identified alterations in taxa corresponding to disease status (8 ASVs), age (30 ASVs), and race (9 ASVs; see Supplemental Table 3). These included several taxa that have been previously reported to discriminate between healthy and atopic individuals including *B. vulgatus* (33) (ASV1, ASV230)*, Prevotella copri* (ASV8, ASV50) (34, 35), and a *Ruminococcus* species (36) (ASV123; Figure 1D).

**Figure 1:**
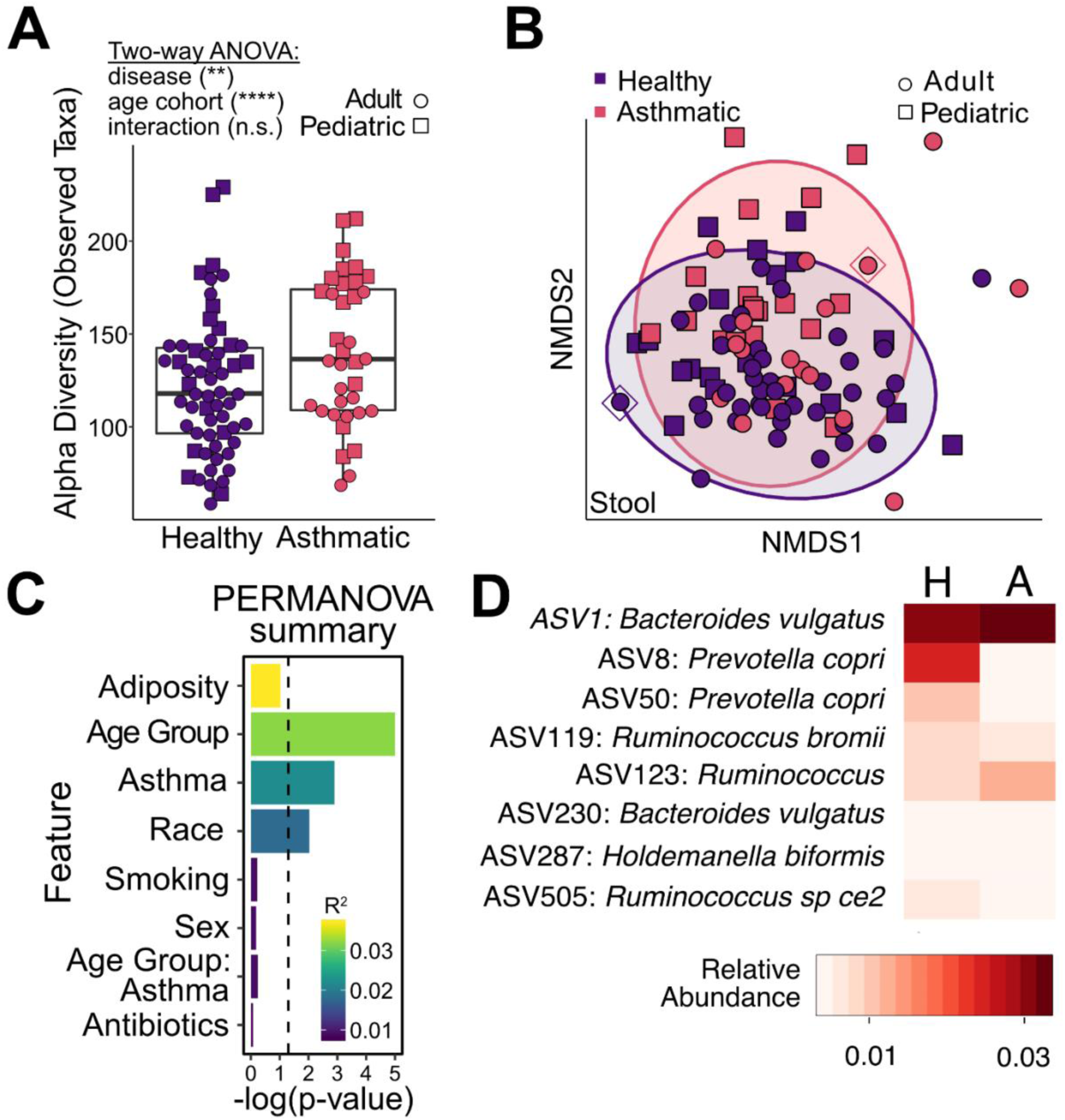
V4-16S rRNA profiling of stool from MARS cohort identifies gut microbiome differences in patients with asthma. **A)** Richness based on observed number of ASVs in stool samples from the MARS Cohort. **B)** Non-metric multidimensional scaling (NMDS) on Bray-Curtis dissimilarity of MARS gut microbiomes. Ellipses represent 95% confidence intervals; diamonds indicate donor dyad (MARS0022/MARS0043). **C)** Bar plot summarizing results of PERMANOVA analysis on the beta diversity in (B). We also tested the homogeneity of the variance using PERMDISP2 and found no detectable difference in dispersion associated with age group or asthma status. Color represents R^2^ value and the length of the bar represents −log(p-value). Dashed line indicates a p*-*value threshold of 0.05. **D)** Heatmap of average relative abundances of differentially abundant taxa identified between patients with asthma and healthy controls using DESEQ2.

We next sought to test whether the asthmatic gut microbiota could affect pulmonary inflammation in a mouse model of AAI. To explore this question, we selected a pair (“dyad”) of human fecal microbiota samples from a healthy and asthmatic subject to “humanize” gnotobiotic mice by oral gavage. We initially selected an unrelated healthy-asthma dyad where the individuals were demographically similar (Subjects 0043 and 0022; matched for age, sex, BMI, and smoking history, see Supplemental Table 1). We constructed a Naïve Bayes Classifier (NBC) to generate several metrics that would help us evaluate the suitability of the selected dyad for characterization in a gnotobiotic animal model (see Methods for additional details). First, to quantify how similar each sample was to its respective cohort, we calculated a Sample Score that ranged from 0 (typical of healthy) to 1 (typical of asthma) and found that both candidate donor microbiomes were typical of their respective disease cohorts (Figure 2A and Supplemental Figure 2). Second, we also visualized samples by Feature Score (Figure 2B, see Methods) to confirm our selected samples cluster with their respective cohorts. Third, to evaluate the testable microbial relationships in the selected dyad relative to all other possible selections, we counted the number of ASVs, for all possible dyads agnostic to host demographics, whose relative abundance was consistent with the NBC’s learned differences between the asthma and healthy cohorts. We defined this metric as a Pairwise Feature Score (PFS) which is greater than zero for every taxon in a dyad whose relative abundances are concordant with our model’s predictions (see Methods and Supplemental Figure 3). We compared the number of “model concordant taxa” (PFS>0) between all possible dyads (Figure 2C, Supplemental Figure 4C) and found that our selected dyad contained a greater than average proportion of model concordant taxa. This indicates that the number of testable microbial comparisons mirroring the relationship between the asthma and healthy cohorts within our selected dyad is typical among all dyads. Together, these results support the idea that the demographically well-matched MARS0022-0043 dyad is characteristic of the microbial differences between cohorts and showcases a new tool for characterizing microbiome dyads.

**Figure 2:**
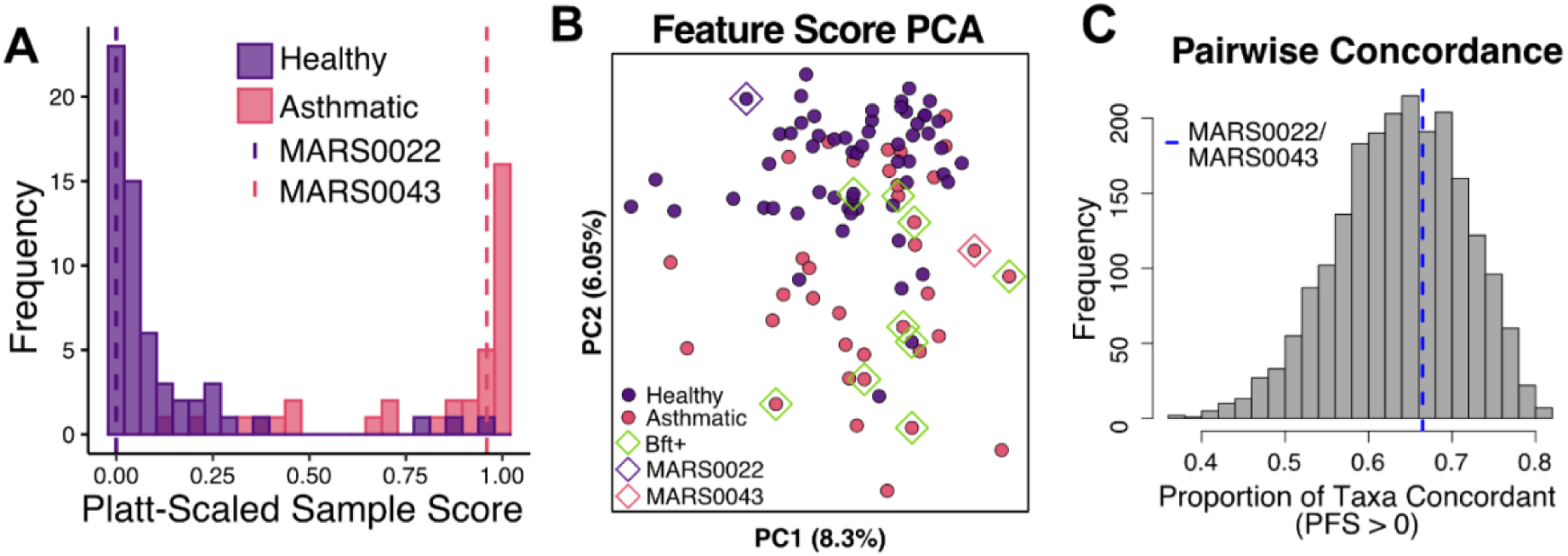
Selected donors capture discriminatory microbial relationships. **A)** Histogram of Platt-scaled MARS NBC Sample Scores. A score of 0 is consistent with a “healthy appearing” sample and a score of 1 is consistent with an “asthma appearing” sample. Vertical dashed lines denote the sample scores for MARS0022 (purple) and MARS0043 (pink). **B)** PCA of NBC Feature Scores, calculated as the log likelihood that the relative abundance for a given taxon would occur in the healthy or asthma cohorts. MARS0022 (purple diamond) and MARS0043 (pink diamond) are the donor samples used in subsequent experiments in gnotobiotic mice. Green diamonds denote *bft* positive samples. **C)** Histogram of the proportion of pairwise concordant taxa across all possible healthy-asthma donor dyads. Vertical dashed line denotes the MARS0022 and MARS0043 dyad.

Mice humanized with the “healthy” (originating from MARS0022) or “asthmatic” (MARS0043) fecal microbiota underwent ovalbumin sensitization and challenge (OSC) or were sacrificed after 4 weeks without OSC, resulting in four groups of mice (abbreviated as depicted in Figure 3A). To gain an overview of the effect of the gut microbiota on lung inflammation we performed bulk RNA-Seq on whole lungs. As expected, mice undergoing OSC had transcriptome profiles that were distinct from naïve mice, reflecting upregulation of genes and pathways related to Type 2 and eosinophilic inflammation demonstrating that we successfully induced AAI in HO and AO mice (Supplemental Figure 5 Supplemental Table 4, 5).

**Figure 3:**
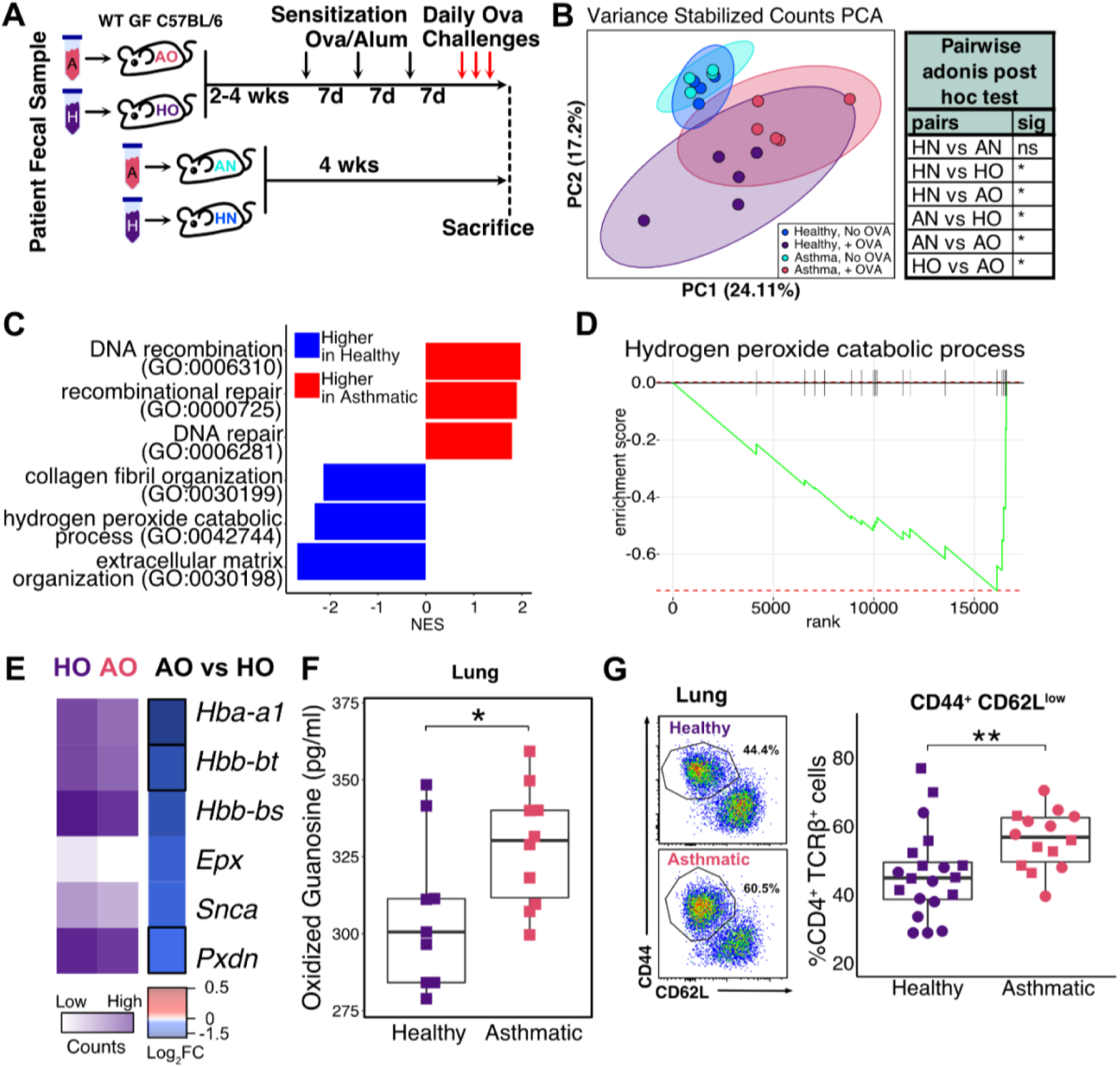
An asthmatic microbiota alters allergic airway inflammation phenotype in humanized gnotobiotic mice. **A)** Overview of experimental paradigm. O: recipients of ovalbumin sensitization and challenge (OSC); N: no ovalbumin sensitization or challenge; A: recipient of fecal sample from human donor with asthma; H: recipient of healthy donor fecal sample. **B)** Principal Components Analysis of variance-stabilized gene counts from lung RNA-Seq (n=4,5). (PERMANOVA, 999 permutations: OSC (p = 0.001), donor (p = 0.022), and donor:OSC interaction (p=0.037); Results of a post-hoc test are also shown in the table. **C)** Bar plot showing normalized enrichment score (NES) of select GO pathways upregulated (red) or down-regulated (blue) in AO mice compared to HO mice (n=5) (p-adjusted<0.05). **D)** GSEA enrichment plot of the hydrogen peroxide catabolic process pathway upregulated in the lungs of AO mice compared to HO mice. **E)** Heatmap of individual hydrogen peroxide catabolic process genes in AO and HO mice. In purple scale: log-transformed size factor normalized counts. In blue-red scale: Log-2 fold change of AO compared to HO. Genes outlined in black represent a p < 0.05. **F)** Oxidized guanosine in AO and HO mouse lungs (n=9-10 mice per group; Wilcoxon, one-tailed). **G)** Flow cytometry of the lungs of AO and HO mice comparing effector T-cells (CD44+CD62L^low^, CD4+TCRb^+^) from the lungs of AO and HO mice (n=6-10 mice/group; 2 experiments denoted by shape).

We next evaluated the effect of the gut microbiota on lung transcriptome profiles. In naïve mice, the gut microbiota appeared to have little or no effect on the overall transcriptome profiles since the difference between HN and AN mice was negligible (Figure 3B). In contrast, mice undergoing OSC colonized with different microbiota had distinct lung transcriptome profiles (see HO vs. AO mice in Figure 3B), suggesting the gut microbiota may be important in the context of AAI. Notably, expected allergy-related genes (*Il4, Il5, Il13*) and pathways (Type 2 immune response, hyperreactivity, eosinophil, and neutrophil pathways) that were upregulated by the induction of AAI were not differentially expressed between HO and AO mice (Supplementary Figure 5B). Further, we found no difference in the degree of sensitization to OVA or expression of Th2 related cytokines (*Il4*, *Il5*, or *Il13*) in OSC treated mice (Supplemental Figure 5A). These data suggest that Th2-related responses do not make up the transcriptomic differences between HO and AO mice.

Differentially abundant pathways between HO and AO mice included several involved in DNA repair and recombination (Figure 3C, Supplemental Figure 6, Supplemental Table 4, 5). We also saw enrichment of a pathway associated with the breakdown of hydrogen peroxide (GO:0042744) in HO compared to AO mice (Figure 3C, D). Many of the genes in this pathway have been implicated in protective responses to oxidative stress. These include hemoglobin synthesis genes (*Hba-a1, Hbb-bt,* and *Hbb-bs*), which are known to be upregulated and protective during oxidative stress in extra-erythropoietic tissues (37, 38), and peroxidasin (*Pxdn*), an enzyme which is likewise known to play a protective role during oxidative conditions in tissues (Figure 3E) (39). When considered with the known roles of oxidative stress in upregulating DNA damage and repair machinery in AAI (40, 41), we hypothesized that increased oxidative stress led to DNA damage in AO mice. To test this idea, we measured oxidized guanosine, a marker for oxidative stress (42), in the lungs of mice and found that it was increased in AO compared to HO groups (Figure 3F). Together, these data reflect increased oxidative stress in the lungs of mice that received the asthmatic microbiota in the context of airway inflammation.

We further investigated the impact of the microbiota on immune cell subsets by performing immunophenotyping on tissues from HO and AO mice. Flow cytometry of lung tissue demonstrated an increase in effector T cells in AO mice compared to HO mice (TCRβ^+^CD4^+^CD62LloCD44^+^ cells; Figure 3G). Consistent with RNA-Seq results, we did not observe a difference in neutrophils or eosinophils recovered from the lung tissue (Supplemental Figure 7C, D). However, we observed an increase in CD4+ T cells expressing IL-17A after restimulation and intracellular staining (Supplemental Figure 7E). Coupled with the increased transcription of *Il17a* in the lung tissues (Supplemental Figure 7B), these findings are consistent with enhanced Th17 cell recruitment to the lungs of AO mice compared to HO mice. This response appeared to be localized to the lungs, since we did not observe increased Th17 cells in the spleen, nor increased serum IL-17A protein (Supplemental Figure 7E,F). Taken together, these results support the idea that the gut microbiota did not change the expression of allergy-associated pathways but did demonstrate alterations in Th17 responses and increased markers of oxidative stress in the lungs of OSC treated mice.

We performed V4-16S rRNA profiling of humanized gnotobiotic mice to better understand how gut microbial community ecology influences immune responses in the lung. This analysis affirmed that the gut microbial communities present in the gnotobiotic mice strongly resembled the human donors from which they had originated (Supplemental Figure 8A-C). While human fecal transplantation into gnotobiotic mice can result in some reconfigurations of the original community (43), our analysis showed that many pairwise concordant taxa identified by our NBC colonized AO and HO gnotobiotic mice. These taxa included *B. uniformis*, *B. fragilis*, and Erysipelotrichaceae (*Longicatena caecimuris*), of which the latter two have been previously implicated in asthma pathogenesis (44–46) (Supplemental Figure 8D).

We next used IgA-seq to identify microbes that affect gut mucosal immune responses in gnotobiotic mice. Our IgA-seq analysis showed that *Bacteroides* species were more likely to be coated by IgA in AO mice compared to HO mice (Figure 4A, B). In AO mice, these IgA-coated bacteria included *B. uniformis* and most prominently, *B. fragilis*. We isolated and sequenced the *B. fragilis* strain found in the asthmatic microbiota, referred to here as BFM04319, and found that its genome encoded for the *B. fragilis* toxin gene, *bft*, also called fragilysin.

**Figure 4:**
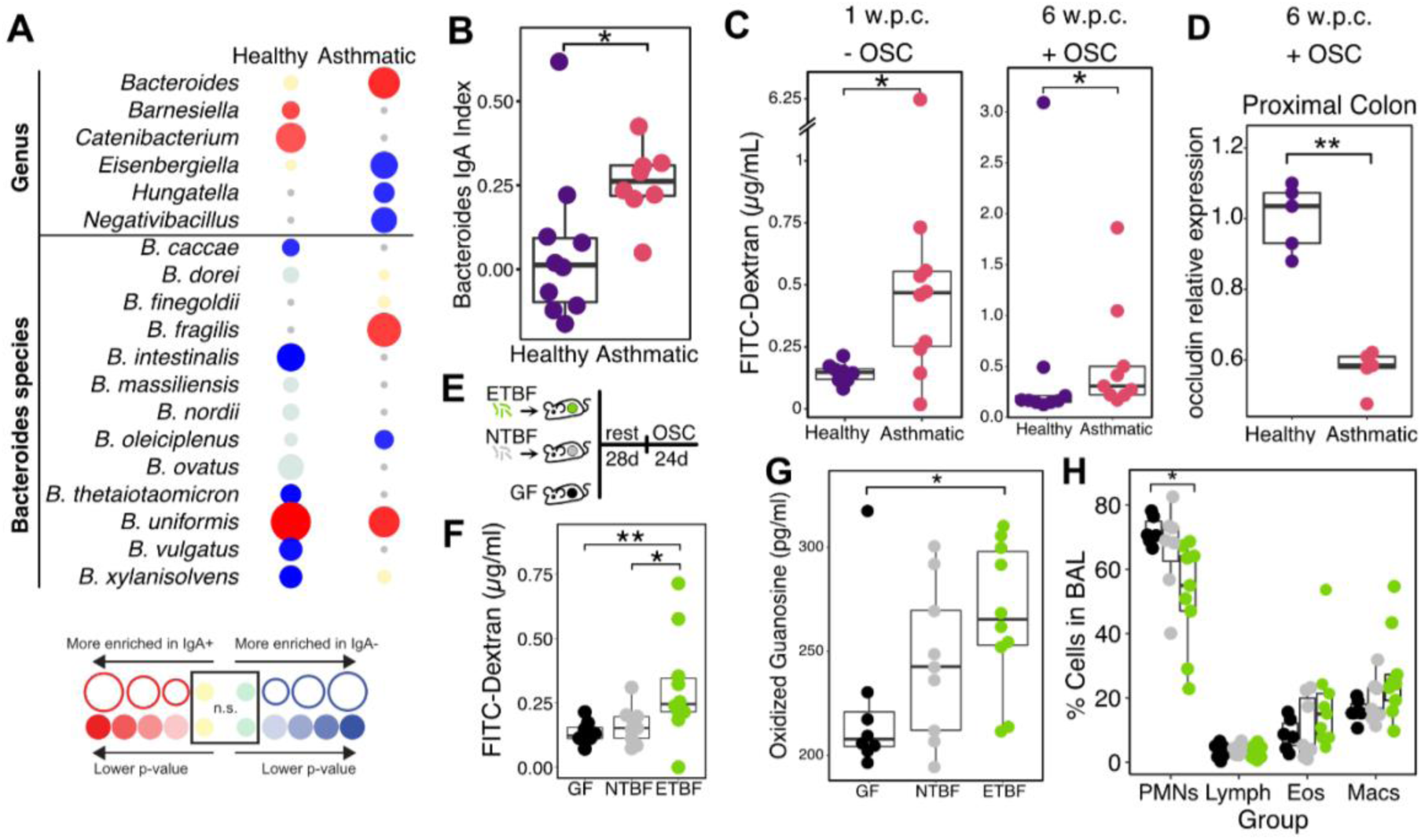
Enterotoxigenic *Bacteroides fragilis* from human donor with asthma alters gut barrier permeability in humanized gnotobiotic mice. **A)** Bubble plot of IgA-Seq results from AO and HO mice at the genus level (n=8-10 mice/group), and specific species of *Bacteroides.* Bubble color indicates significant enrichment (red) or depletion (blue) in the IgA coated fraction. Bubble size indicates the magnitude of enrichment as determined by the IgA index. **B)** IgA Index of Bacteroides genus between AO mice and HO mice colonized with asthmatic or healthy microbiota (n=8,10 mice/group). **C)** Intestinal permeability of mice colonized with asthmatic and healthy microbiota following a 1-week and 6-week colonization (n=8-10 mice/group). **D)** qPCR of occludin gene (*Ocln*) in the proximal colon of humanized mice (n=5 mice/group). **E)** Overview of the experimental paradigm testing the ability of *bft* carrying enterotoxigenic *B. fragilis* (ETBF) and *bft* lacking non-toxigenic *B. fragilis* (NTBF) to affect gut barrier function in mice. **F)** Intestinal permeability of mice colonized with either ETBF or NTBF and germ-free controls (GF) (n=8-10 mice/group; Kruskal-Wallis, one-tailed Dunn post hoc with Benjamini-Hochberg correction). **G)** Oxidized guanosine in lungs of ETBF, NTBF, and GF mice (n=8-10 mice/group; Kruskal-Wallis, one-tailed Dunn post hoc with Benjamini-Hochberg correction). **H)** Cytospin cell counts from bronchoalveolar lavage collected from mice colonized with ETBF, NTBF or germ-free controls. PMN=polymorphonuclear cells, Lymph=lymphocytes, Eos=eosinophils, Mac=macrophages. (n=8-10 mice/group).

Fragilysin is a *B. fragilis* diarrheal toxin (47) whose proteolytic activity is directed against the adherens junction protein E-cadherin and disrupts the intestinal epithelial barrier (48, 49). We reasoned that gut barrier function would be impaired in AO mice so we assessed FITC-Dextran leakage in the humanized gnotobiotic mice and found an increase in intestinal permeability in mice colonized with the asthmatic microbiota starting as early as 1 week after gavage and persisting after OSC (Figure 4C). Similar to previous studies (50), increased FITC-Dextran leakage was also accompanied by downregulation of the gene encoding for occludin (*ocln*), an important protein involved in the regulation of intestinal tight junctions, in the colons of AO mice (Figure 4D).

Since gut barrier dysfunction has been previously implicated in inducing systemic oxidative stress (50, 51), we hypothesized that ETBF in our asthmatic stool sample was responsible for the increased oxidative stress in the lungs of AO mice by disrupting the gut barrier. We directly tested whether ETBF alone are sufficient to modulate oxidative stress after OSC by examining three groups of gnotobiotic mice: 1) mice that remained germ-free (GF); 2) mice monocolonized with BFM04319 (ETBF), and 3) mice monocolonized with a non-toxigenic *B. fragilis* strain VPI2553 (NTBF) (Figure 4E) (52). As expected, mice colonized with ETBF had greater intestinal permeability than either GF controls or NTBF colonized mice (Figure 4F). We observed higher levels of oxidized guanosine in the lungs of ETBF colonized mice compared to GF controls and no statistically significant difference between NTBF colonized and GF mice (Figure 4G). While not statistically significant, oxidized guanosine levels in ETBF colonized mice trended higher than in the NTBF colonized mice (Figure 4G). Further, mice colonized with ETBF had lower neutrophil counts in their bronchoalveolar lavage compared to NTBF-colonized mice or GF controls, although eosinophils, lymphocytes, alveolar macrophages and *Il17* transcription were not significantly different (Figure 4H, Supplemental Figure 7G). These results provide evidence that colonization with ETBF is sufficient to cause gut barrier dysfunction, modulate airway inflammatory profile, and increase oxidative stress during following OSC.

Following our identification of ETBF as an important driver of oxidative stress in the lungs of gnotobiotic mice undergoing OSC, we then sought to investigate the importance of ETBF in the context of different microbial communities. While exploring multiple donors does not guarantee translatability to humans (43), it does reveal the strength and robustness of the ETBF-oxidative stress phenotype. Therefore, we selected multiple dyads made up of one ETBF+ donor with asthma and one ETBF- healthy donor for humanization and measured gut barrier permeability, lung cytokine gene expression, and pulmonary oxidative stress after OSC. To select donor samples, we screened MARS stool samples for *bft* using PCR and qPCR (53, 54) and identified five asthmatic fecal samples harboring ETBF and five healthy fecal samples lacking ETBF, each pair matched by both age group and community composition metrics derived from our NBC (see Methods; Supplemental Table 1). Using the selected communities, we humanized germ-free mice and performed OSC. We evaluated humanization in recipient gnotobiotic mice by comparing the Bray-Curtis dissimilarity scores from 16S rRNA sequencing of stool from human donors and recipient mice at the time of sacrifice. This led us to exclude three groups of humanized mice (2 ETBF-, 1 ETBF+) whose engrafted fecal microbiome did not best reflect their donors, leaving seven humanized mouse groups (Figure 5A, Supplemental Figure 9A-C; see Methods).

**Figure 5:**
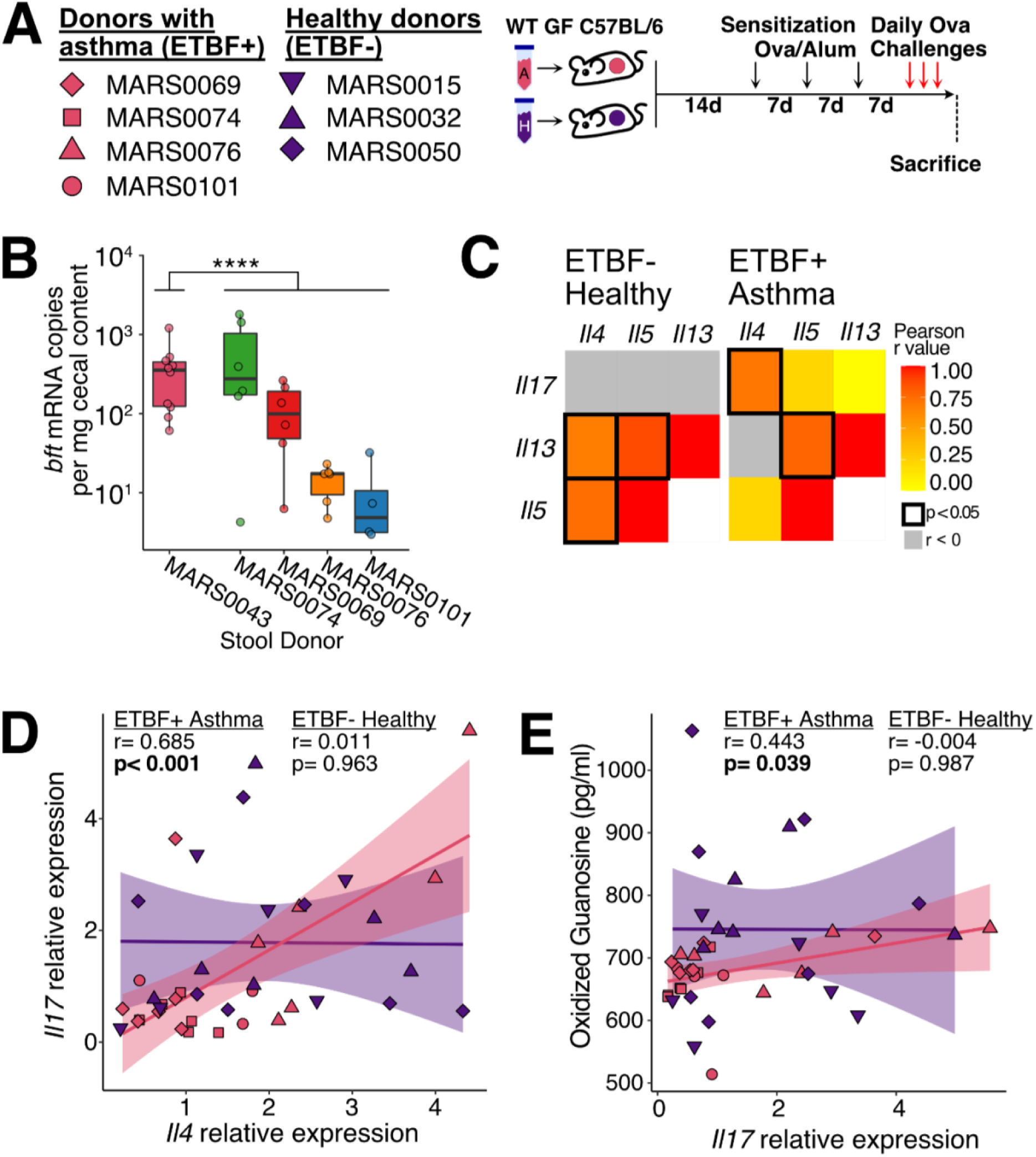
Humanization with multiple microbiota suggests ETBF produced *bft* drives a phenotype in pulmonary inflammation. **A)** Overview of experimental paradigm. Five healthy and five asthmatic microbiota were used to humanize germ-free mice (4 - 7 mice per donor). Two weeks later, pulmonary inflammation was induced by OSC. After evaluating humanization, only seven mouse groups were used in subsequent analyses. **B)** qPCR measurements of *bft* in the cecal content of humanized mice (t-test against initial MARS0043 microbiome). **C)** Correlations between RT-qPCR measures of lung cytokines from humanized mice in healthy microbiota and asthmatic microbiota. **D)** Correlation between RT-qPCR measures of *Il4* and *Il17* in the lungs of humanized mice (Stieger’s test for the difference between two independent correlations p = 0.04). **E)** Correlation between oxidized guanosine in the lungs of humanized mice and RT-qPCR measures of lung *Il17*.

Overall, we found no differences in gut barrier leakage, pulmonary oxidative stress, or gene expression of lung cytokines between mice colonized with an asthmatic ETBF+ microbiota and those receiving an ETBF- healthy microbiota (Supplemental Figure 9D-F). To understand the differences between these experiments and our original donor experiment, we measured copies of *bft* RNA and DNA in the cecal contents of ETBF+ humanized mice and found that the median expression levels of the mice with newly selected human samples were 1.2- to 67.5-fold lower than that of the original donor sample, MARS0043 (Figure 5B, t-test p=0.01). These results are consistent with previous research showing that ETBF vary their enterotoxin expression based on the microbial community context (55) and may explain the lack of an ETBF- oxidative stress phenotype seen in these mice.

Given that the expression of *bft* was far more variable than observed in the original donor dyad, we tested if there was any correlation between *bft* expression and markers Th17 or AAI. We observed that despite the variance in *bft* expression, ETBF+ asthmatic humanized mice had an altered pattern of cytokine expression within the lung compared to ETBF- healthy humanized mice (Figure 5C). As expected, the expression of Th2 cytokine-encoding genes *Il5* and *Il13* were highly correlated to each other in both experimental groups. However, we found that *Il4* expression was highly correlated with *Il17a* expression in mice colonized with ETBF+ asthmatic communities (Figure 5D; Pearson = 0.685, p < 0.001), whereas the correlation between *Il4* and *Il17* expression in mice colonized with ETBF- healthy communities was significantly weaker (Pearson = 0.011, p = 0.963; Stieger’s test for the difference between two independent correlations p = 0.01). Previous studies have identified a correlation between *Il17* expression and oxidative stress in the lungs (56). We found that ETBF+ humanized mice displayed the expected correlation between lung *Il17a* expression and oxidative stress (Figure 5E, Pearson = 0.443, p = 0.039), but the ETBF- healthy humanized mice did not (Pearson = −0.004, p = .987). Together, these results suggest that ETBF+ microbiota from donors with asthma may link AAI to *Il17a*.

Finally, we investigated the importance of *bft* in humans by examining its prevalence in the stool of MARS participants. Using PCR and qPCR (53, 54), we screened all human fecal specimens containing *B. fragilis* by 16S rRNA sequencing for *bft* and found a total of 8 individuals with asthma and 2 healthy subjects with detectable *bft*, indicating a significant enrichment of *bft* in subjects with asthma compared to healthy individuals (Figure 6A and Figure 2B). We then asked whether higher rates of ETBF could be a consequence of an increased prevalence of *B. fragilis* in the asthma cohort but found no difference in the frequency of *B. fragilis* colonization (Figure 6B). Even amongst only individuals colonized with *B. fragilis*, patients with asthma were still more likely to be colonized with ETBF than NTBF compared to healthy individuals (Figure 6C). We next examined fecal calprotectin levels from remaining subjects to determine whether or not asthma was associated with gut barrier permeability. Overall, we found that calprotectin was either low or undetectable in all samples but more likely to be detectable in samples originating from a donor with asthma (Fisher’s test p-value = 0.053, Figure 6D). While the low levels of calprotectin are likely due to long-term sample storage, these results tentatively suggest a degree of barrier dysfunction in patients with asthma.

**Figure 6:**
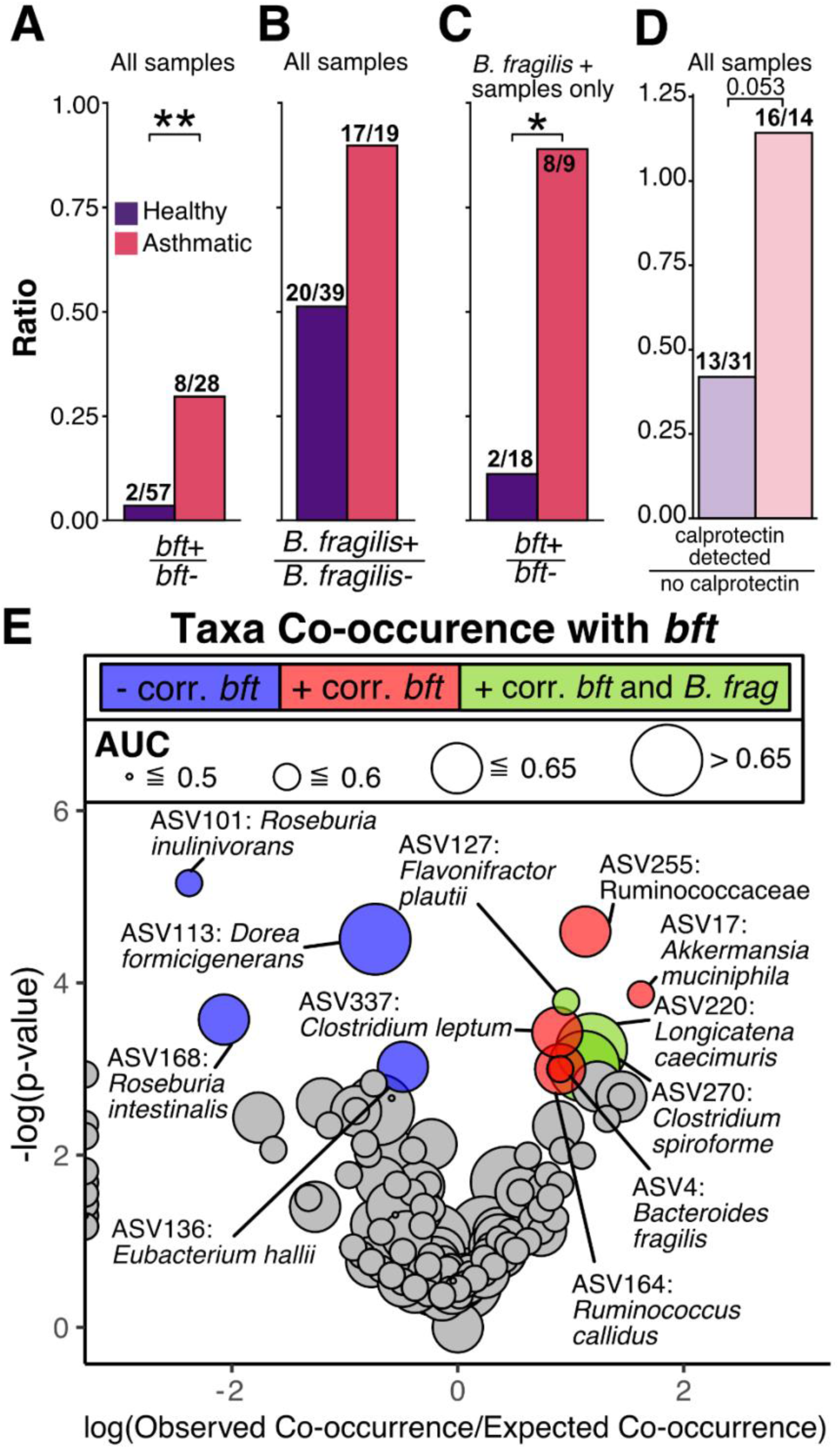
*bft* is enriched in individuals with asthma. **A-D)** Ratios of **(A)** *bft* positive over *bft* negative samples (n=59 Healthy, 36 Asthmatic). *bft* positive samples were characterized by PCR and qPCR screening among the 37 total subjects with *B. fragilis*. The 58 remaining subjects without *B. fragilis* by 16S rRNA sequencing were not screened and were considered *bft* negative. **B)** *B. fragilis* positive samples over *B. fragilis* negative samples, characterized by V4 16S rRNA community profiling (n=59 Healthy, 36 Asthmatic). **C)** *bft* positive samples with *B. fragilis* over *bft* negative samples with *B. fragilis* (n=20 Healthy, 17 Asthmatic). **D)** Stool samples with detectable calprotectin via ELISA over those with no detectable calprotectin. **E)** Volcano plot describing co-occurrence between *bft* and taxa in human stool samples, calculated as the log ratio of the number of samples in which *bft* was observed in the same sample as the taxon compared to the number of times *bft* and the taxon would be expected to occur in the same sample by chance alone^73^. Color represents a significant positive (red) or negative (blue) co-occurrence with *bft* after multiple hypothesis correction. Taxa positively correlated with *bft* and *B. fragilis* (ASV4) are shown in green. Gray represents a non-significant relationship. Size represents the area under the ROC curve (AUC) of the taxa calculated from the NBC. (n=59 Healthy, 36 Asthmatic)

We suspected that other taxa in the gut may be associated with ETBF, so we performed a co-occurrence analysis with *bft* presence and our 16S rRNA sequencing results (Figure 6E). In addition to *B. fragilis*, we identified 7 taxa positively correlated and 4 negatively correlated with ETBF colonization. Among the positively correlated taxa, were 4 taxa co-occurred with *bft* but not *B. fragilis* (ASV4), suggesting that ETBF may influence gut microbiota composition in a different manner from NTBF species. Notably, *bft* co-occurs with *Dorea formicigenerans* (AUC = 0.663, Rank 11), *Longicatena caecimuris* (AUC = 0.662, Rank 12), and *Clostridium spiroforme* (AUC = 0.657, Rank 13), which are highly discriminatory between asthma and healthy individuals according to our NBC (Supplemental Figure 2D), implying that the presence of *bft* reinforces asthma-associated changes in the gut microbiota.

## Discussion

Despite our growing understanding of the origins of asthma, the heterogeneous nature of the disease remains a barrier to treatment. In addition to factors such as age (57), gender (58), smoking status (59), and microbial exposures (60), the gut microbiota is increasingly appreciated as a determinant of asthma risk (9, 45). Here we leverage a cross-sectional human clinical study of 95 patients with and without asthma and use humanized gnotobiotic mice to show that, in the right community context, enterotoxigenic *Bacteroides fragilis* (ETBF) can increase Th17 inflammation and oxidative stress in the lungs of mice with AAI, potentially by disrupting the gut barrier. While our focus on established disease precludes us from exploring the early microbial origins of asthma, our experiments identified a potential microbial influencer of ongoing pathophysiology.

Humanization of gnotobiotic mice is a powerful tool to study the human microbiota and its effects on host phenotypes, but has many well-established caveats. First, the human microbiota does not perfectly maintain community and functional structure between donors and mouse recipients (43, 61). Second, virulence factors adapted to human hosts may not affect mice to the same degree. Third, a gut microbiota may influence the host phenotype by multiple mechanisms simultaneously, such as in asthma where multiple microbial metabolites are known to alter the course of the disease, each by their own respective mechanisms (9, 11–13). Together, these issues can complicate the interpretation of results from gnotobiotic experiments and affect the translatability of those results back to humans.

Designing experiments to address these limitations requires the selection of donor samples that both control for confounding factors that modify the microbiome apart from the disease of interest as well as capture the relevant microbial relationships between healthy and disease-affected populations. Control of confounding factors is often accomplished by matching the clinical demographics of donors, which shape the composition of the gut (43, 62) (Figure 1C). However, there is no standardized practice for identifying samples that capture the most microbial relationships relevant to disease. Given the immense interpersonal variability of the microbiome (63–65), no single pair of samples captures all the discriminatory microbial features from a population but we can still select pairs enriched in these relationships to study. Here we devised a Bayesian model that quantified the representative nature of each of our samples and cataloged the important population-wide differences in community composition captured by any dyad. We believe that using such an approach for sample selection will be useful in identifying microbial drivers of disease in future clinical studies.

Our in-depth profiling of a single dyad revealed an ETBF+ microbiome from a donor with asthma increased the Th17 response and oxidative stress within the lungs of humanized gnotobiotic mice in the context of AAI. While previous studies have implicated airway microbes in inducing Th17 responses in AAI (24, 66), our work provides evidence that this phenotype can be mediated by a gut microbiota-expressed factor. In follow-up monocolonization experiments, we confirmed that the ETBF isolated from the donor with asthma causes increased gut barrier permeability and pulmonary oxidative stress in the context of pre-existing inflammation. Notably, we did not measure an increased Th17 response in the lungs of ETBF monocolonized mice compared to NTBF or GF mice. This could mean that other community members in addition to ETBF are important for inducing systemic (67) and lung Th17 responses.

We also explored the effect of community context on ETBF by colonizing groups of mice with ETBF+ communities from multiple donors with asthma. We did not detect increased gut permeability, pulmonary oxidative stress, or lung Th17 responses in the ETBF+ humanized mice. However, we did observe that overall expression of *bft* was much lower in this experiment compared to the MARS0043. Despite this, ETBF+ asthmatic microbiota demonstrated a link between *Il17a* and *Il4*, suggesting that ETBF may mediate an association between Th17 responses and AAI. Taken with our monocolonization data, these results imply that *bft* may influence inflammation in the lungs, but that its expression and impact on asthma may be modulated by community context. Based on these results, we propose that *bft* and ETBF may contribute to the clinical heterogeneity of asthma in humans.

While gnotobiotic experiments offer control over many features of the gut microbiota, they ultimately remain a proxy of asthma in humans. Confirming the findings from our humanized mouse models will require a longitudinal human clinical study including ETBF colonized subjects with asthma and controls that undergo rigorous phenotypic characterization that incorporates measuring gut barrier permeability, circulating microbial products, and immunophenotyping of the blood and bronchial lavage. The results of this study would establish whether ETBF could adversely influence asthma in people and define the environmental and microbial circumstances that enable ETBF to cause these phenotypes. Additionally, the outcomes of such a clinical study could also link two previously disparate features of asthma: first, that increased gut permeability (68, 69) and increased gastrointestinal symptoms (70) have been observed in patients with asthma, and secondly, that increased oxidative stress is associated with more severe asthma with a higher rate of exacerbations and corticosteroid resistance (2, 3).

We acknowledge that ETBF is unlikely to be a universal mechanism contributing to asthma and would probably only apply to a subset of people living with asthma (22% from our study). On the other hand, implicating a barrier-disrupting organism in asthma pathology could lead to unique interventions to improve asthma control. For instance, targeting ETBF colonization by vaccination, antibiotics, or phage therapy could offer a novel means of manipulating the immune manifestations of asthma. Alternatively, candidate therapeutics designed to improve gut barrier function through their effects on tight junctions for other diseases including arthritis (71) and celiac disease (72) could be repurposed to modify the gut-lung axis in asthma. Unlike early childhood interventions that aim to alter the development of asthma, these therapies may provide benefit to patients with established disease.

## Methods

### The Microbiome and Asthma Research Study

The Microbiome and Asthma Research Study (MARS) was designed to investigate the role of intra- and interpersonal variation in the gut microbiome on human asthma pathogenesis. This cohort has been described in a previous manuscript (24). Briefly, all individuals were either healthy or had moderate-to-severe asthma and had not received oral corticosteroids or antibiotics in the 30 days prior to enrollment. Individuals were included in the asthma cohort if they had at least one positive skin prick test or serum aeroallergen-specific IgE. Two age cohorts were recruited representing a pediatric (aged 6-10 years) and an adult (aged 18-40 years) population. Of 104 patients initially enrolled in the study, 103 were asked to provide samples and fill out the questionnaire. Of the 103 asked, 6 were excluded due to insufficient medical documentation and 2 did not provide stool samples (Supplemental Figure 10). As summarized in Supplemental Table 1, a total of 95 patients provided stool and relevant demographic data, including antibiotic and asthma medication use over the past year, and were ultimately included in this study. Fecal collection tubes with spoons and toilet hats were provided to patients after enrollment. Patients either provided a fecal specimen at the recruitment visit or collected and stored a fecal sample at home at −20° C for no more than 24 hours before returning the sample to the study site where it was stored at −80° C until processing.

### Experimental Animals

Germ-free C57BL/6 mice were bred and maintained in sterile flexible vinyl isolators. Sterility was assured by monthly monitoring of mouse stools by 16S rRNA gene PCR amplification as well as aerobic and anaerobic culture. Germ-free mice were maintained on a strict 12 hour light cycle and a diet of autoclaved mouse chow (LabDiet: Standard Diet 5021 - Autoclavable Mouse Breeder). For each experiment no more than five mice were housed in a cage. Mice were randomly assigned to experimental groups, with the exception of age and gender matching mice in each experimental group. Investigators were not blinded to experimental groups. Male and female mice within a group were caged separately but housed and handled within the same flexible vinyl isolator.

### Isolation, Growth, and Whole Genome Sequencing of *Bacteroides fragilis* strains

*B. fragilis* strain BFM04319 was isolated from MARS0043 (subject with asthma) stool using anaerobic culture methods. Briefly, a 10 mg/mL stock of homogenized stool in 10% glycerol/PBS was thawed in a Coy anaerobic chamber, plated on BHI/mucin agar containing 0.1% mucin, 0.1% Resazurin, and 0.05% L-Cysteine (HCl) and 1.2 mg/L histidine/hematin and incubated at 37°C for 6 days. Single colonies were isolated in BHI/mucin liquid media and stocked in 10% glycerol. Culture purity and identity was confirmed by V4 16S sequencing. Non-toxigenic *B. fragilis* VPI2553 was provided as a generous gift from Dr. Jeffrey I. Gordon. All strains were grown anaerobically in BHI/mucin media as above at 37°C for 24 hours before diluting 1:2 with 20% glycerol, freezing, and gavaging into GF mice.

Genomic DNA was extracted from *B. fragilis* strain BFM04319 by phenol/chloroform extraction. We then used an adaptation of the Nextera Library Prep kit (Illumina, cat. FC-121-1030/1031) (73) and sequenced on a MiSeq to achieve ∼80X coverage of the 5Mbp ETBF genome. Reads were trimmed by quality and adapter content with bbtools (sourceforge.net/projects/bbmap/). Scaffolds were created with SPAdes (74) and annotated with prokka (75). Our assembly had an N50 of 432688 and an L50 of 5. BFM04319 had an average nucleotide identity of 98.84% to the genome of the *B. fragilis* type strain VPI2553 (NCBI reference sequence: CR626927.1) (76).

### Processing of Stool and 16S rRNA Profiling Library Preparation

Human stool was pulverized in a biosafety cabinet with liquid nitrogen using a pestle and mortar and aliquoted into 50 - 100 mg samples and stored at −80° C prior to use. For both human and mouse fecal specimens, crude DNA was extracted using phenol:chloroform:isoamyl alcohol and homogenized with a bead beater using sterilized zirconium and steel beads as previously described (77). The aqueous layer was then purified with a 96-well QIAGEN PCR Clean up kit and quantitated by measuring the absorbance at 260/280 nm. DNA concentrations were normalized to 5 ng/μL and 10 ng of DNA was used to PCR amplify the V4 16S rRNA region using barcoded primers as previously described (78). PCR-amplified DNA was pooled to equal concentration and the library purified using AMPure XP SPRI beads. DNA was sequenced using a MiSeq with 2×250 bp chemistry. All samples had a minimum read-depth of 5000.

### Analysis of 16S rRNA data

Fastq files were demultiplexed and binned into amplicon sequence variants (ASVs; Supplemental Table 6) using DADA2 as previously described (24). Taxonomic determination of ASV sequences to the lowest possible level was performed with RDP Classifier (79) using a database built to permit species level identification (77) with a minimum bootstrap support of 80%.

ASVs were normalized using total sum scaling. Diversity analysis of 16S data was carried out with vegan (v2.5-7) and phyloseq (v1.28.0) in R. Richness was estimated as the average count of observed taxa after rarefying to 5000 reads using the vegan rarefy function. PERMANOVA was carried out for 100,000 iterations to achieve a minimum p-value of 10^-5^ using adonis2 in vegan. PERMDISP2 was performed with the betadisper function in vegan. Differential abundance analysis was carried out with DESEQ2 (version 1.24.0) as previously described (80, 81). Batch effect in 16S data resulting from sequencing run differences was assessed using PERMANOVA and found to contribute significantly to the variance in the data, but did not otherwise affect our results (see Supplemental Table 2). Co-occurrence was calculated using the cooccur package (82).

### IgA-Seq

We performed IgA-seq on mouse fecal samples as previously described (77). In brief, fecal samples were prepared in reduced PBS, stained with polyclonal goat anti-human IgA (Abcam #ab96998) or goat anti-mouse IgA (Abcam #ab97104) fluorescently labeled with DyLight649. Following antibody staining, samples were washed with PBS and resuspended in 100 mM HEPES and 150 mM NaCl containing a 1:4000 dilution of Syto-BC (Invitrogen). Samples were run and acquired on a BD Aria II maintained in a laminar flow biosafety cabinet. Input, IgA- and IgA+ positive fractions were acquired and sequenced as previously described (83). To analyze these data we used either the IgA index (77) or a Bayesian method to predict the probability of a bacteria being found in either the IgA+ or IgA- fractions (84).

### Mixture Distribution Naïve Bayes’ Classifier for 16S Profiling of Asthma vs. Healthy Patients

#### Input Data

Raw ASV counts data were normalized to total counts (relative abundance). To be included in the NBC, a taxon had to appear in at least 7 samples. This number was determined by calculating how many samples are required to identify enrichment of a taxa in either the healthy or asthmatic cohort with 95% confidence, based on presence/absence alone (Binomial Test). In total, 392 ASVs between 95 human stool subjects were included in this analysis.

#### Algorithm

By Bayes’ theorem, the probability that a microbiome composed of many taxa belongs to an individual with asthma is

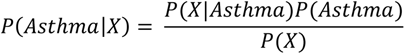

where *P*(*X*|*Asthma*) is the likelihood of a microbiome sample (*X*) occurring given all the microbiome data from the asthmatic cohort, *P*(*Asthma*) is the prior probability of any MARS patient having asthma, and *P*(*X*) is the probability of the microbiome data (*X*) occurring given the entire MARS microbiome dataset (*X*). To accomodate the sparsity and zero-inflation inherent to microbiome data, we built our NBC to fit relative abundance data to a mixture distribution. We model *P*(*X*|*Asthma*) and *P*(*X*|*Healthy*) (see Figure 2A: pink and purple, respectively) as mixtures of (1) a beta distribution of relative abundance when the taxon is present and (2) a binary distribution when the taxon is not detected,

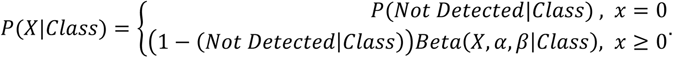

By modeling the frequency of non-detection separately from relative abundance we increase the sensitivity of our model to learn differences in sparse taxa. The prior probability of a patient having asthma (*P*(*A*)) was determined based on the proportion of patients with asthma used in the training data. The beta distribution for each taxon was fit by Maximum a Posteriori estimation using Newton’s Method, given a Gaussian prior. Hyperparameters of the prior distributions were optimized by a grid search. Using mixture distributions generated for each taxon in the model, we constructed a Naïve Bayes’ Classifier in R to predict patient asthma status based on microbiome composition. The NBC produces metrics that are useful for further analysis, including a Feature Score which can be described as the log likelihood ratio of a taxon occurring at a given relative abundance in stool from an asthma donor compared to from a healthy donor. The Feature Score per taxon (*i*) per sample (*j*) is 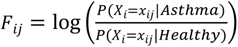.

For each sample, the Feature Scores for all taxa (*n*) in a sample are summed to calculate the Sample Score, which can be described as the log likelihood of the sample being from the asthma population rather than the healthy. The Sample Score per sample is

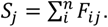

Samples were classified as asthmatic if the Sample Score was positive, and healthy if negative.

We identified taxa concordant with the model in any given healthy-asthma dyad by calculating the Pairwise Feature Score between the two samples for each ASV. The Pairwise Feature Score for a taxon in a dyad is then calculated as the difference between the Feature Score for the taxon in the sample with asthma and the Feature Score of the same taxon in the healthy control is

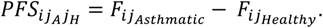

ASVs in a dyad with a positive pairwise feature score were considered concordant with the model. See examples in Supplemental Figure 4.

#### Random Forest

The random forest model (RF) was created in R using the randomForest package (85) (v4.6-14). All forests included 1000 trees (ntrees=1000) with 30×30 tree sampling with replacement (sampsize=c(30,30)) and were built on the same 392 ASVs used in the NBC.

#### Model Evaluation

All AUC and ROC curve values were calculated in R using pROC (v1.16.2). ROC curve and AUC values in Figure 2C were calculated based on the sample scores for the NBC and tree classification votes for the RF. Platt-scaling was performed using the glm (family = binomial(logit)) and predict functions in R. We performed leave-one-out cross-validation (LOOCV) and repeated this process 100 times to estimate an average LOOCV classification rate. The NBC achieved a leave-one-out cross validation (LOOCV) accuracy of 75.8% (Supplemental Figure 2B,C), which is similar to studies of comparable size (22, 86), and performed as well as a Random Forest classifier (LOOCV accuracy: 75.1%), another tool commonly used to classify disease-associated microbiome data. Taxa identified as highly discriminatory by the NBC were highly correlated with those found to be important by Random Forest (Supplemental Figure 2D, Supplemental Table 7, rho = 0.4439, p-value < 0.0001, Spearman Correlation).

### Humanized Gnotobiotic Mouse Model

We began with frozen pulverized fecal samples brought into an anaerobic chamber and transferred into reduced 1X PBS supplemented with 0.1% Resazurin and 0.05% L-Cysteine (HCl) to a concentration of 10 mg/mL. Fecal samples were then thoroughly homogenized using a sterilized probe homogenizer. Resuspended fecal samples were diluted 1:2 in 20% glycerol in PBS/Cysteine and stored in sealed HPLC vials at −80° C until use. Sample viability was confirmed by outgrowth of 100 uL of prepped gavage material on BHI/mucin agar containing 0.1% mucin, 0.1% Resazurin, and 0.05% L-Cysteine (HCl) and 1.2 mg/L histidine/hematin. A minimum of 1×10^4^ CFU/mg was used for colonization. Germ-free mice were humanized by oral gavage of 200 mL of a thawed homogenized stool sample. The microbiota was allowed to stabilize over two to four weeks prior to further intervention. All experiments were performed with at least one asthmatic and one healthy microbiota and separate donor groups were maintained in separate isolators to prevent cross-contamination. For the additional humanization experiments seen in Figure 5, we selected asthmatic microbiota containing ETBF, of which 5 were viable in culture outgrowth. For each donor with asthma, we selected a healthy sample within the same age group – either adult or pediatric – that did not contain ETBF but had the highest pairwise feature score (Supplemental Figure 9B). We assessed the quality of human microbiota transplantation by calculating the Bray-Curtis dissimilarity of 16S rRNA sequencing data between human donors and murine recipients. We then ranked the average dissimilarity between recipients and donor and only included recipient groups who were the top ranked in similarity to their human donor.

Allergic airway inflammation was induced in mice using chicken egg ovalbumin as previously described (24, 87). Germ free mice were sensitized on days 0, 7, and 14 by intraperitoneal injections of 200 μL of OVA/alum: ovalbumin (50 μg, Sigma grade V) combined with Imject Alum (Thermo Scientific) as per the manufacturer’s recommendations. Mice were challenged on days 20 - 22 by intranasal introduction of 1 mg ovalbumin suspended in 50 μL sterile PBS while under anesthesia. Control mice were neither sensitized nor challenged unless otherwise noted.

### Immune Cell Isolation from Tissues

Cells were extracted from tissues as described (24). Briefly, lungs were minced and incubated in digestion buffer (0.2 U/ml Liberase DL (Roche Applied Sciences) and 0.2 mg/ml DNase (Sigma) in Hank’s Buffered salt Solution (without Ca^2+^/Mg^2+^) for 25 min at 37°C before being passed through a 70µm cell strainer (88). Spleen and lymph nodes were dissociated manually and passed through a 70 µM cell strainer. Red blood cells were removed from lung and spleen samples by treatment with ACK lysis buffer.

### Flow Cytometry of Isolated Immune Cells

Data were acquired on a FACSCanto II (BD Biosciences) equipped for the detection of eight fluorescent parameters. The following antibodies were used: PE anti-mouse SiglecF (Clone E50-2240; BD Pharmigen™), FITC anti-mouse CD4 (Clone GK1.5, Biolegend), FITC anti-mouse CD11c (Clone N418, Biolegend), PE anti-mouse CD44 (Clone IM7, BD Pharmigen™), PE anti-mouse IL-17A (Clone TC11-18H0.1, Biolegend), PerCP-Cy™5.5 anti-mouse TCR β chain (Clone H57-597, BD Pharmigen™), PE-Cy™7 anti-mouse CD11b (Clone M1/70, BD Pharmigen™), PE/Cyanine7 anti-mouse CD62L (Clone MEL-14, Biolegend), APC anti-mouse Ly6G (Clone 1A8, Biolegend), APC anti-mouse CD45 (Clone 30-F11, Biolegend), PerCP anti-mouse CD45 (Clone 30-F11, Biolegend), APC/Cyanine7 anti-mouse I-A/I-E (Clone M5/114.15.2, Biolegend), eFluor450 anti-mouse FoxP3 (Clone FJK-16s, eBioscience™), eFluor450 anti-mouse IL-13 (Clone 13A, eBioscience™), Brilliant Violet 421™ anti-mouse F4/80 (Clone BM8, Biolegend), APC/Cyanine7 anti-mouse TCR β chain (Clone H57-597, Biolegend), PE/Cyanine7 anti-mouse IFNγ (Clone XMG1.2, Biolegend), APC anti-mouse TNFα (Clone MP6-XT22, BD Pharmigen™). Intracellular staining of cytokines was conducted as previously described (24). Briefly, cells were stimulated for 4 h at 37°C with PMA (10ng/mL), ionomycin (200ng/mL), monensin (1:1000), and brefeldin A (1:1000). LIVE/DEAD Fixable Aqua Dead Cell Stain Kit was used to assess cell viability in all panels. Data analysis was performed using FlowJo version 10 or higher software (Treestar, Ashland, OR). Gating strategies are summarized in Supplemental Figure 11.

### Transcriptional Profiling of Mouse Lungs

We isolated total RNA from mouse lungs and performed transcriptomic analysis as previously described (24). Reads were mapped to the mouse genome using bowtie2 (v2.3.4.1) (89), quantified at the gene level using htseq (v 0.9.1) (90), and differentially expressed genes were identified using DESeq2 (v1.24.0) (81). Functional pathways altered during colonization and/or OSC were identified using gene set enrichment analysis FGSEA (fgsea R package; v1.10.1) (91) with KEGG and GO databases. PERMANOVA was performed using adonis (vegan R package; v2.5-7) and the post hoc test was performed with pairwise.adonis (pairwiseAdonis, v0.3).

### Lung RT-qPCR

RNA from flash-frozen, pulverized lung tissue or lung tissue stored in RNALater (Invitrogen cat. AM7021) was extracted by probe homogenization in TRIzol Reagent followed by chloroform phase separation. DNase I (Qiagen) was then used to degrade DNA according to the manufacturer and this reaction was carried onto Qiagen’s RNeasy Mini kit for purification of RNA. We then quantified the RNA with a Quanti-iT Ribogreen RNA Assay kit (Invitrogen cat. R11490) and synthesized cDNA with a high-capacity RNA-to-cDNA kit (AB cat. 4387406). RT-qPCR was performed on a Biorad CFX96 Real-Time System using Power SYBR Green PCR Master Mix (AB 4367659). Primer pairs used in this paper are shown in Supplemental Table 8 including GAPDH as the reference gene. Samples were run in triplicate and were excluded if the range of raw Ct values for target or reference exceeded 2.

### Protein Quantification

Mouse Serum IgE specific to ovalbumin was quantified by sandwich ELISA (24). Briefly, plates were coated with 10 μg/mL purified ovalbumin overnight at 4° C and then blocked with PBS 1x with 1% BSA. Sera were diluted 1:10 and plated alongside purified OVA-specific IgE (Clone 2C6, AbD Serotec) as a standard curve. The plate was incubated for 2 hours at room temperature. Bound IgE was detected using goat anti-mouse IgE-HRP (Clone RME-1, Biolegend). Serum IL-17A was measured as part of a LEGENDPlex Multiplex protein assay (Biolegend) following the manufacturer’s protocol.

### DNA/RNA Oxidative Damage ELISA

To estimate pulmonary oxidative stress we measured oxidized guanosine (8-hydroxyguanosine, 8-hydroxy-2’-deoxyguanosine, and 8-hydroxyguanine) in the lungs of mice using an ELISA (Cayman Chemical cat. 589320) following the manufacturer’s instructions and analysis template.

### Intestinal Permeability Assay

In order to assess intestinal permeability, FITC-Dextran gut-to-serum absorption was measured at the time of sacrifice as previously described (92, 93). Briefly, a baseline blood sample (150 - 250 uL) was taken from each mouse via facial vein puncture. After one hour of fasting, the mice were orally gavaged with 200 μL of 40 mg/ml FITC-Dextran (4 kDa; Sigma Aldrich) in sterile PBS. Food and water were then withheld for an additional 45 minutes. After three hours mice were sacrificed and blood was collected by cardiac puncture. Serum was separated from blood samples using serum separator tubes according to manufacturer’s instructions (BD microtainer). Fluorescence of pre- and post-gavage serum were measured (at a 5-fold dilution in PBS) using an excitation wavelength of 485nm and an emission wavelength of 528nm. A standard curve from 0 to 40 μg/ml FITC-Dextran read on the same plate was used to convert the RFU values to concentration of FITC-Dextran. Finally, pre-gavage serum FITC-Dextran concentrations were subtracted from post-gavage serum FITC-Dextran concentrations to quantify leakage from the gastrointestinal tract to the circulation.

### Cytospin

Bronchoalveolar lavage was collected by flushing the mouse lungs using 1 mL of 0.1% BSA in sterile PBS. 5×10^5^ cells of each sample were loaded onto a slide using a Shandon Cytospin 2. Slides were then methanol fixed and stained with eosin and methylene blue following kit directions (ThermoScientific Shandon KwikDiff Stains). Cells were counted using a bright-field microscope at 400X magnification. 300 non-red blood cells were counted per sample, with careful scanning to ensure no repetition between high power fields.

### Screening for *bft* in Subject Stool Samples

Purified DNA from stool samples containing *Bacteroides fragilis* based on V4-16S rRNA community profiling were normalized to 5 ng/μL. PCR was used to amplify the constant c-terminal region of the *bft* gene (Supplemental Table 8) (53). The results of the reaction were assessed using a 2% agarose gel stained with GelRed (Biotium). Previous literature has identified this method suffers from low sensitivity, so we also conducted qPCR on the same samples (54). qPCR primers were designed based on the *bft* sequence identified in BFM04319 and verified by confirming amplification against purified BFM04319 genome, and a complete fecal community harboring BFM04319 (AO stool gDNA), but not in a community lacking BFM04319 (HO stool gDNA). A single band of the expected size was detected by electrophoresis of BFM04319 genome amplification and AO stool gDNA amplification, but not HO stool gDNA. A dilution series of *B. fragilis* genome was used to demonstrate primers were sufficiently sensitive to detect *bft* from less than 2*10^-5^ ng of *B. fragilis* genome. The presence of a band or a Cq value of less than 35 was considered a positive result and indicated the presence of *bft* in patient stool.

### Quantitation of *bft* expression in mouse cecal contents

Mouse cecal content RNA and DNA was extracted using the QIAgen AllPrep PowerFecal DNA/RNA kit as per kit instructions (cat. 80244). Nucleic acids were quantitated and cDNA synthesis was performed on 450 ng of cecal RNA using the Lambda Biotech EasyScriptPlus cDNA Synthesis kit (cat. G236) and a primer specific to *bft* (see Supplemental Table 8). qPCR was performed as previously described. Copies of *B. fragilis* genome were estimated by comparing cycle number against a dilution series of purified BFM04319 genome and then normalized to the amount of nucleic acid per milligram of cecal content.

### Quantitation of calprotectin in human stool samples

Human fecal calprotectin was measured using the Calprotectin ELISA Assay Kit (Eagle Biosciences cat. CAL35-K01) following manufacturer’s directions. Between 50 and 100 mg of pulverized human stool was used for each assayed sample.

## Data Availability

All 16S rRNA and genome sequencing data available at European Nucleotide Archive (https://www.ebi.ac.uk/ena/browser/home) under project accession number PRJEB45298.

## Code Availability

Code necessary to run the mixture distribution-based NBC is available via github (github.com/naomiwilson/g<u>nominator</u>)

## Statistics

Statistics and analysis were all performed in R Version 3.6.3. Data are presented as mean with SEM. Statistical significance was conducted using an unpaired Wilcoxon test or Kruskal Wallis test with a post hoc Dunn test where appropriate. Adjustment of p-values for multiple hypotheses was performed using Benjamini-Hochberg correction. Boxplots display IQR and whiskers display 1.5*IQR. The following symbols were used to designate significance: *p < 0.05, ** p < 0.01, *** p < 0.001.

## Study Approval

MARS was approved by the Washington University Institutional Review Board (IRB ID# 201412035). Written informed consent documents were obtained from all MARS subjects or their legal guardians. All animal studies conformed to ethics of animal experimentation and were approved by the Institutional Animal Care and Use Committee (IACUC Protocol ID #: 20180286).

## Acknowledgements

We would like to thank our clinical study coordinators Tarisa Mantia, Caitlin O’Shaughnessy, and Shannon Rook; the physicians of Washington University Pediatric and Adolescent Ambulatory Research Consortium, especially Dr. Jane Garbutt; the Volunteer for Health registry; and the MARS participants and their families. We would also like to acknowledge Dr. Devesha Kulkarni, for her technical expertise as well as the Center for Genome Sciences Sequencing Core and Genome Technology Access Center at the McDonnell Genome Institute for sequencing services. The authors also thank Drs. Brain Laidlaw and Leyao Wang for critical reading of the manuscript and Drs Philip Ahern, Neelendu Dey, and Ansel Hsiao for thoughtful feedback on our work. A.L.K. is funded by the AAAAI Foundation Faculty Development Award and the NIH K08 AI113184. A.H-L received funding through the NIH T32 GM007200. J.S-B received funding through the NIH T32 GM007067.

## Author Contributions

N.G.W., A.H.-L., and A.L.K. conceptualized the work. L.B.B. and A.L.K. planned the clinical study. N.G.W., A.H.-L., A.L.R., N.J., R.T.M., J.S.-B., M.A.L., C.P.T. and A.L.K. contributed to the design and conduct of experiments. N.G.W., A.H.-L., A.L.R., N.J., R.T.M., and A.L.K. analyzed the data. N.G.W., A.H.-L., T.R.R., and S.J.S. developed gnominator. N.G.W., A.H.-L., and A.L.K. drafted the manuscript. All authors interpreted the data and contributed to revising the manuscript.

