## Supplementary Figures 1-11 for "The asthma gut microbiota influences lung inflammation in gnotobiotic mice"

### Supplementary Figures and Table Captions

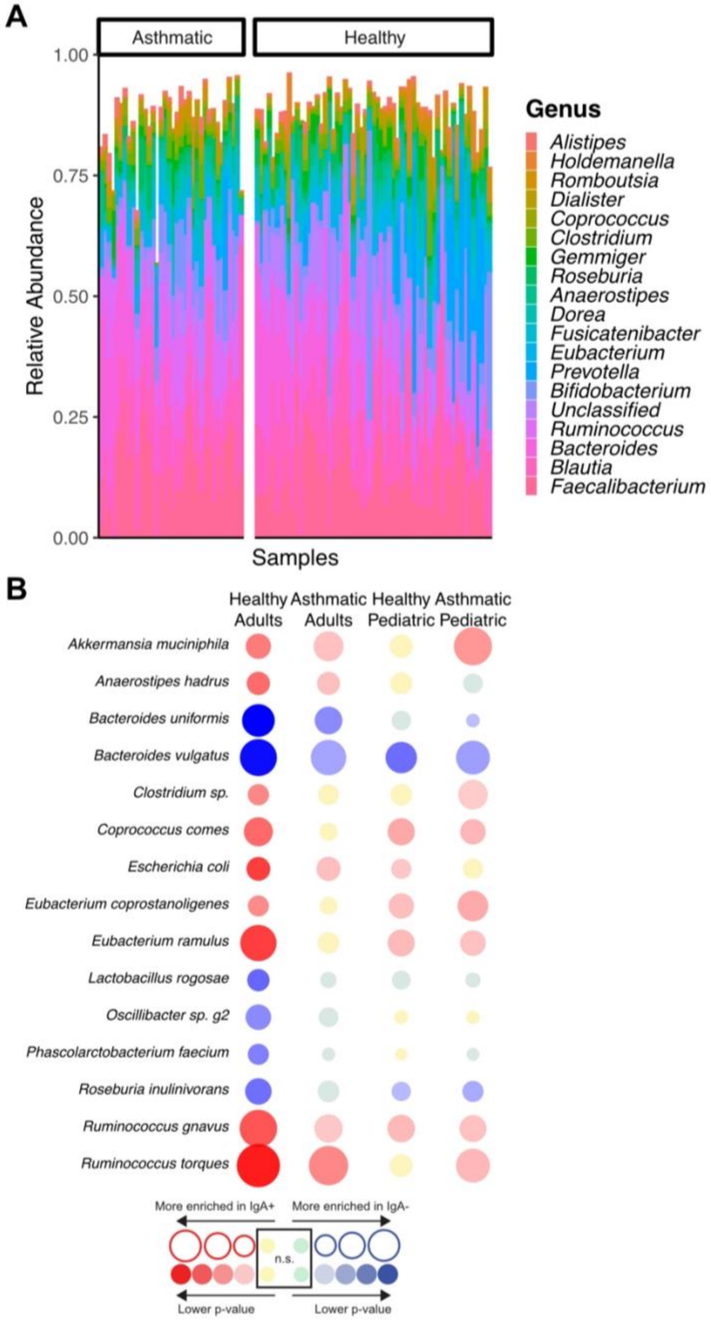


**Supplemental Figure 1: Overview of 16S rRNA sequencing from participants in the MARS A)** Relative abundance bar plot of 95 human fecal samples characterized in this study (n=59 Healthy, n=36 Asthmatic). **B)** IgA coating of bacterial taxa in MARS subject stool was assessed using IgA-Seq. Species-level taxa summarized from ASVs were significantly enriched in the IgA+ or the IgA- fractions in at least one of the four cohort groups. Bubble color indicates significant enrichment (red) or depletion (blue) in the IgA coated fraction. Bubble size indicates the magnitude of enrichment as determined by IgA index.

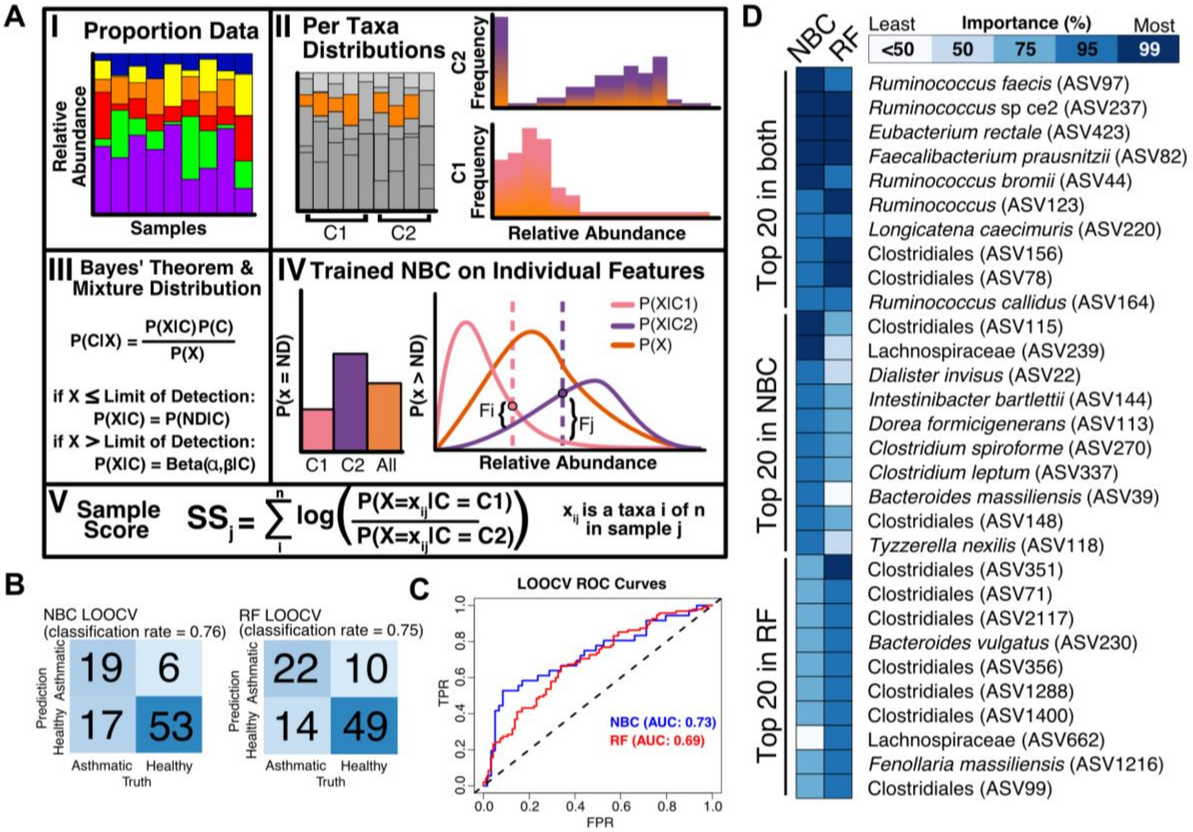


**Supplemental Figure 2: A Naïve Bayes' Classifier is an intuitive tool for the exploration of microbiome data. A)** Overview of Naïve Bayes' Classifier (NBC) fit with a mixture distribution. **I)** Individual counts of taxa by 16S rRNA sequencing are scaled by total read count per sample. **II)** Relative abundances of each taxon across samples are separated by class. **III)** Bayes' theorem can be used to calculate the probability of a given class given the microbiome composition. We model relative abundance as a mixture distribution such that when a taxon is not detected, it is part of a binary distribution and when detected is part of a beta distribution. **IV)** Using our mixture model, an individual taxon can be modeled overall and as part of a class to calculate the probability of belonging to different classes. **V)** The sample score describes the likelihood that a sample is from one class over another. **B)** Confusion matrices summarizing Leave-One-Out Cross Validation (LOOCV) predictions for NBC and Random Forest (RF), respectively. The NBC performed significantly better than chance (p=8.2e-6 by Fisher's exact test). **C)** Receiver Operating Characteristic (ROC) Curves and area under the ROC curve (AUC) scores from LOOCV NBC and RF models. **D)** A comparison of taxa identified as important by NBC and RF. Feature importance in the random forest was inferred based on mean decrease in accuracy. Feature importance in the NBC was inferred based on the AUC of an individual taxa's ROC curve in predicting asthma. Colors correspond to percentiles of ranked importance for taxa considered by each of the models. The top 20 predictive features in each model are presented.

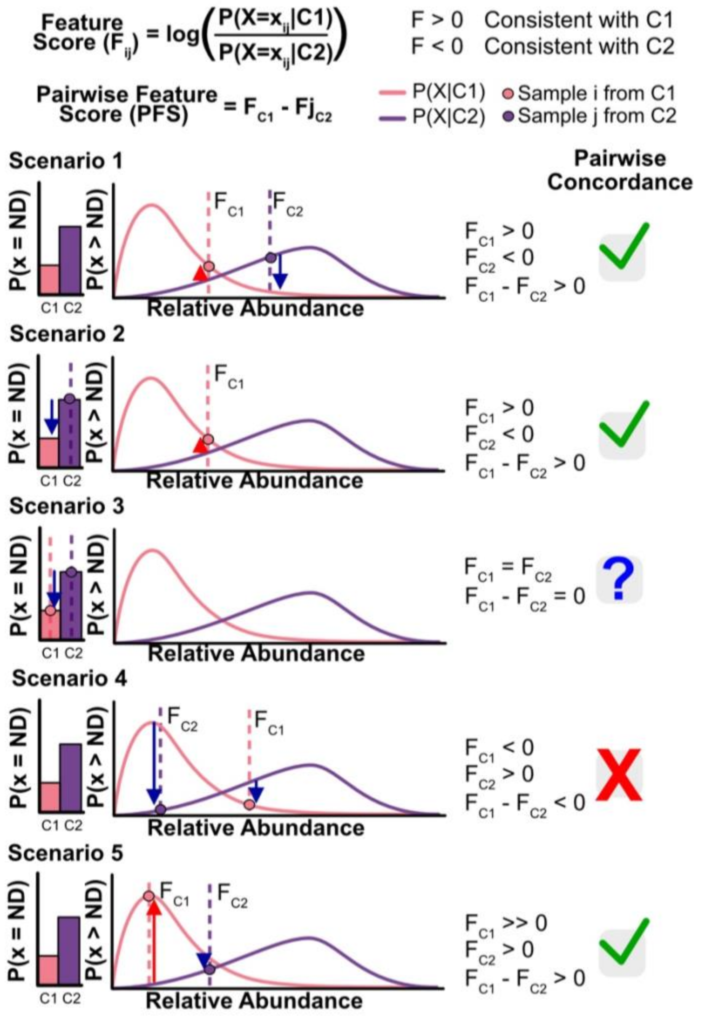


**Supplemental Figure 3: Overview of NBC pairwise concordance metric:** The NBC feature score describes the likelihood that a taxon at a given relative abundance would be found in one class over another. A comparison of the feature scores of the same taxon across two different samples provides the pairwise feature score (PFS*)* that compares the likelihood of these taxa abundances occurring between two samples from different classes. Example scenarios where the pairwise feature score is used to identify concordance with the model are provided. **Scenario 1** shows a case where both the asthma sample and healthy sample are present at relative abundances expected by the model**. Scenario 2** shows a case where a healthy sample is absent, and the asthma sample is present at abundances expected by the model. **Scenario 3** demonstrates a unique situation where the taxon is absent in both samples. In this case, the pair provides no information about the taxon and thus is equal to zero. **Scenario 4** demonstrates the case where the relative abundances of the taxon in both the asthma sample and healthy sample are not consistent with the model. The pairwise feature score in this case is negative, reflecting discordance with the model. **Scenario 5** depicts an edge case, where the relative abundance in the asthma sample is expected, but the relative abundance of the healthy sample is not. The magnitude of the likelihood of the asthma sample is much greater than that of the healthy sample however, and so the pairwise feature score is positive reflecting model concordance in this taxon.

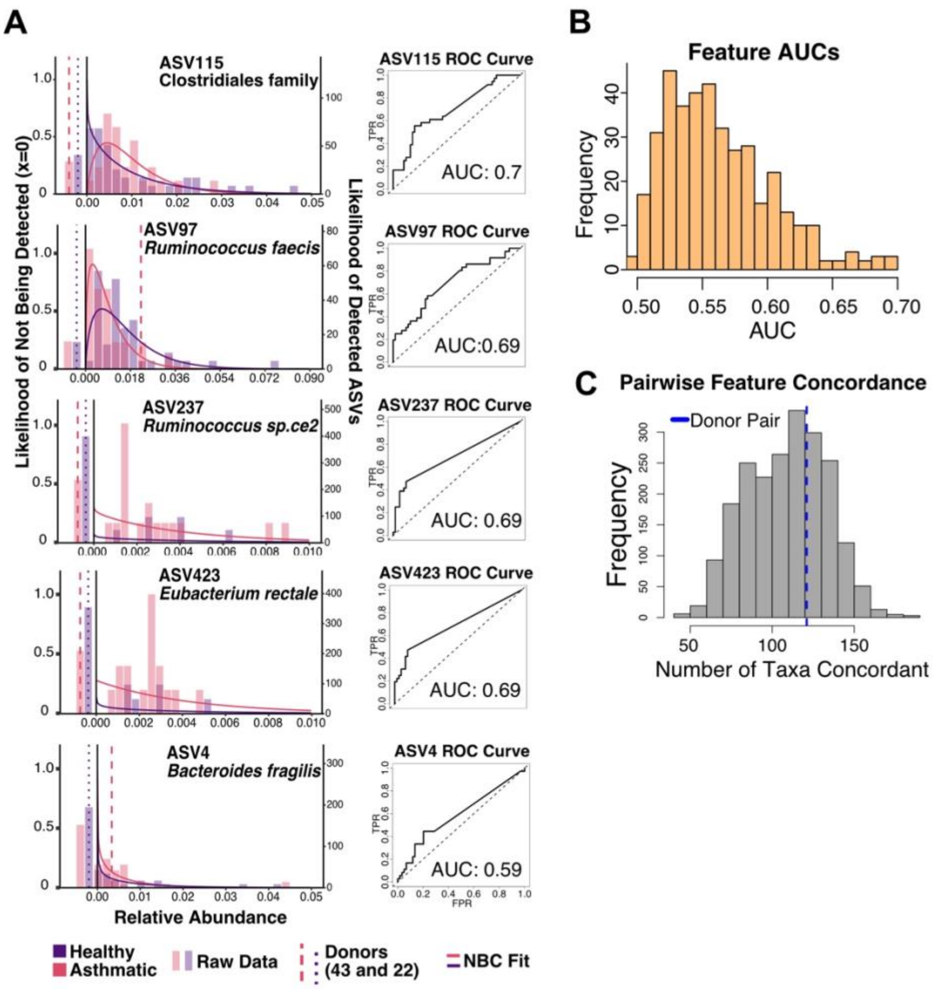


**Supplemental Figure 4: Estimation of taxa importance from mixture distribution and NBC scores A)** Left: raw data histograms overlaid with NBC curve of best fit and right: ROC curve for highly ranked ASVs and *B. fragilis* (ASV4). Donor samples are represented by dashed lines (pink for MARS0043 and purple for MARS0022). **B)** Distribution of AUC values from all 392 ASVs in NBC training set. **C)** Histogram of raw counts of pairwise concordant features per sample. Donor pair (MARS0022/MARS0043) represented by dashed blue line.

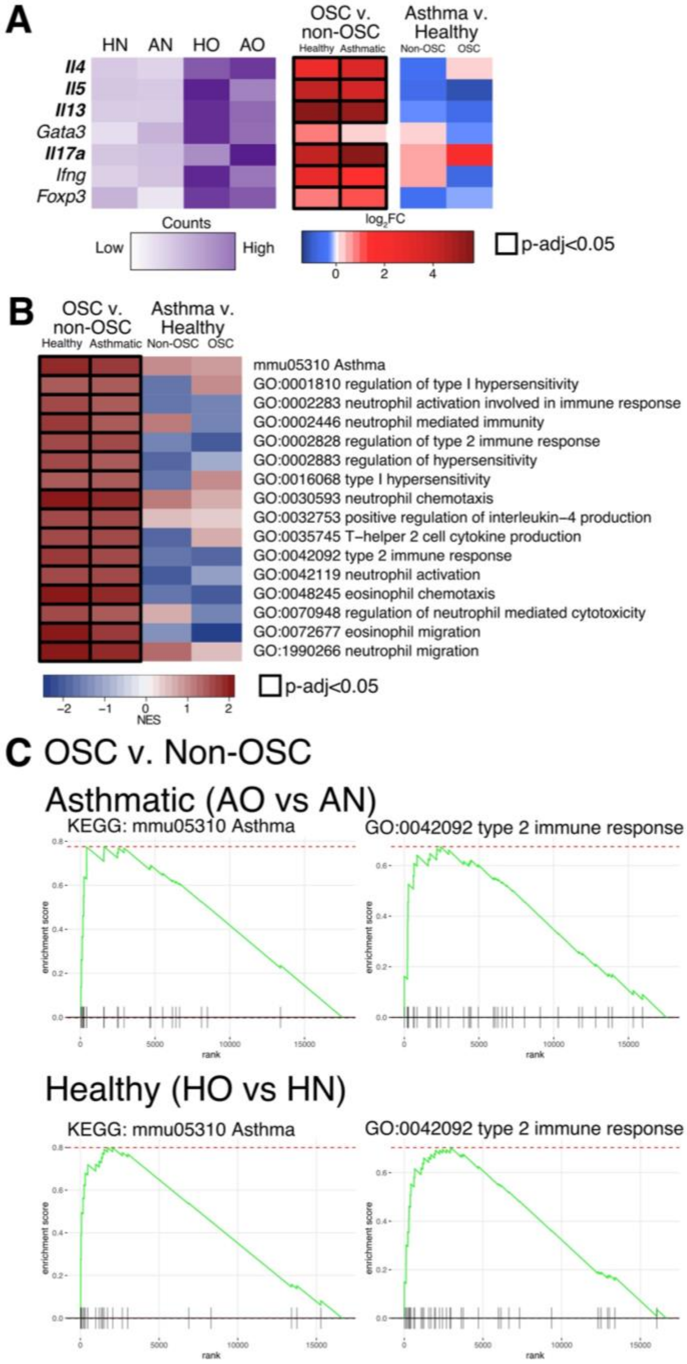


**Supplemental Figure 5: OSC model of AAI induced typical asthma-related and Type 2 immune responses in the lung transcriptome. A)** Heatmap of variance stabilized counts and log2 fold-changes of allergic airway inflammation-related genes. In purple scale: the counts of the genes (z scores scaled by row). In blue-red scale: the log2 fold change. Statistically significant fold changes are outlined in black (n=4-5). **B)** GSEA enrichment heatmap of normalized enrichment scores (NES) of relevant GO and KEGG pathways (n=4-5). **C)** GSEA enrichment plots of the KEGG Asthma pathway and GO Type 2 immune response pathway (n=4).

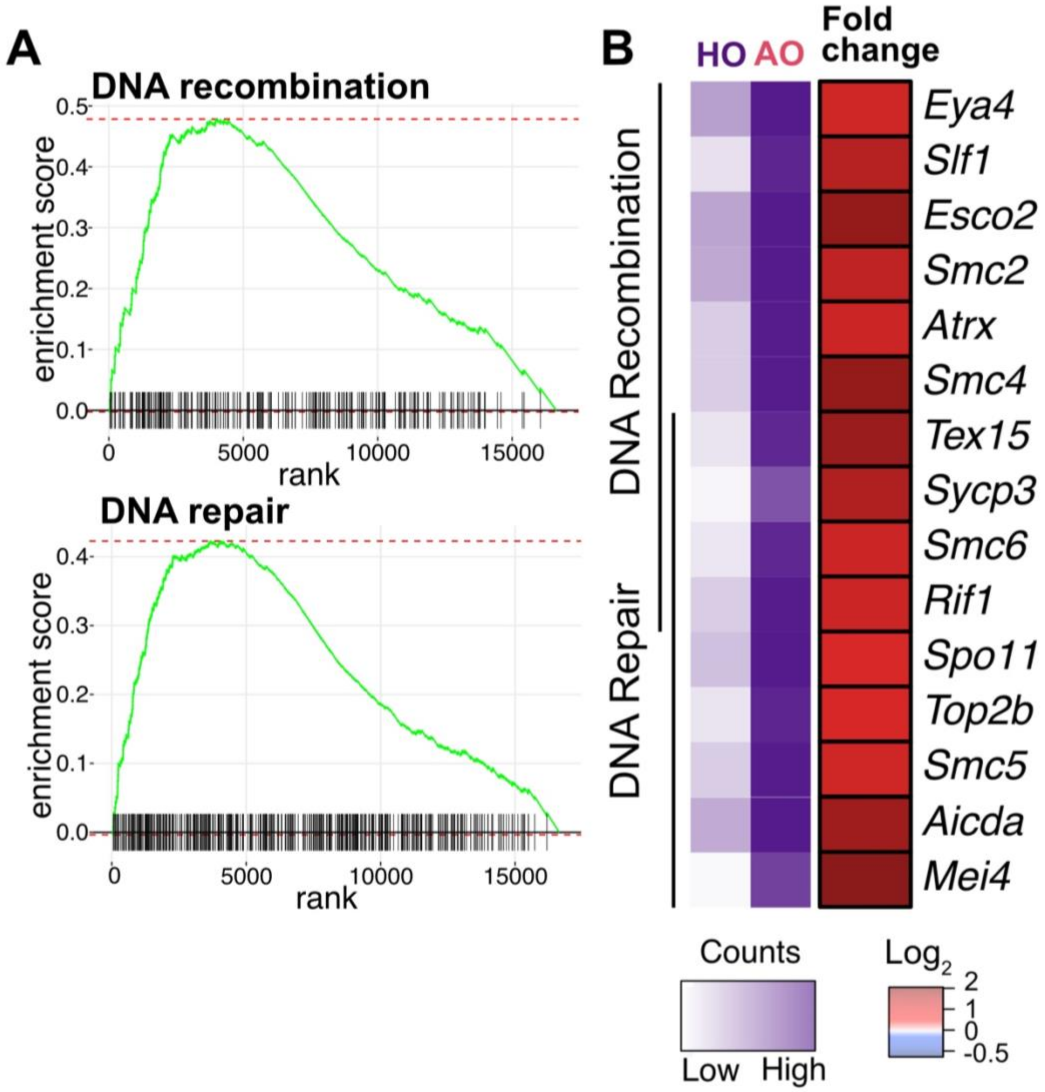


**Supplemental Figure 6: Transcriptomic upregulation of DNA repair and recombination in the lungs of AO mice compared to HO mice A)** GSEA enrichment plots of GO:0006281 DNA repair and GO:0006310 DNA recombination pathways (n=5/group). **B)** Heatmap of variance stabilized counts of leading edge genes from the GO pathways in (A). In purple scale: the counts of the genes (z scores scaled by row). In blue-red scale: the log2 fold change. Statistically significant fold changes are outlined in black (n=5, p-adjusted<0.05).

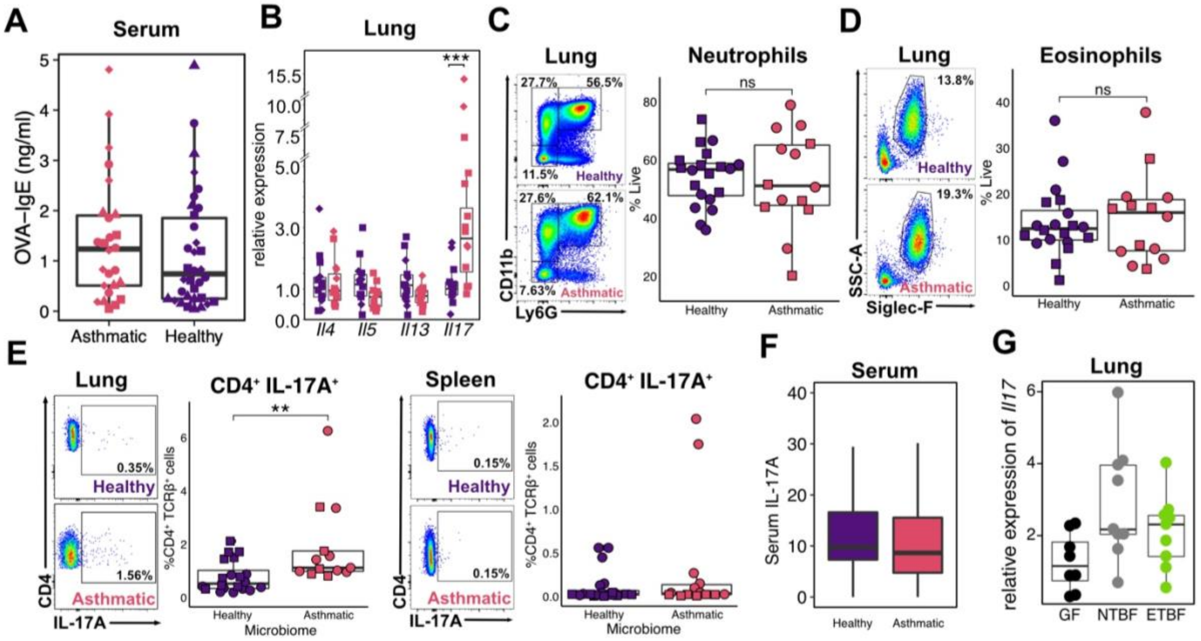


**Supplemental Figure 7: Immunophenotyping of humanized gnotobiotic mice A)** Serum anti-ovalbumin IgE measurements from AO and HO mice (n=6-10/group, 4 experiments denoted by shapes). **B)** Expression of genes encoding Interleukin-4, -5, -13, and -17A measured by RT-qPCR from the lungs of AO and HO mice (n=4-10/group, 2 experiments denoted by shapes). **C - D)** Flow cytometry profiling of granulocytes in the lungs of AO and HO mice (n=6-10/group, 2 experiments denoted by shapes). **E)** Intracellular staining of IL-17A following *in vitro* restimulation from CD4+ T cells isolated from the lungs and spleen of AO and HO mice (n=5-10/group, 2 experiments denoted by shapes). **F)** Serum measurement of IL-17A in AO and HO mice (n=6-10/group, 4 experiments). **G)** Expression of IL-17A measured by RT-qPCR in the lungs of mice mono-colonized with ETBF, NTBF, or germ-free controls (n=8-9).

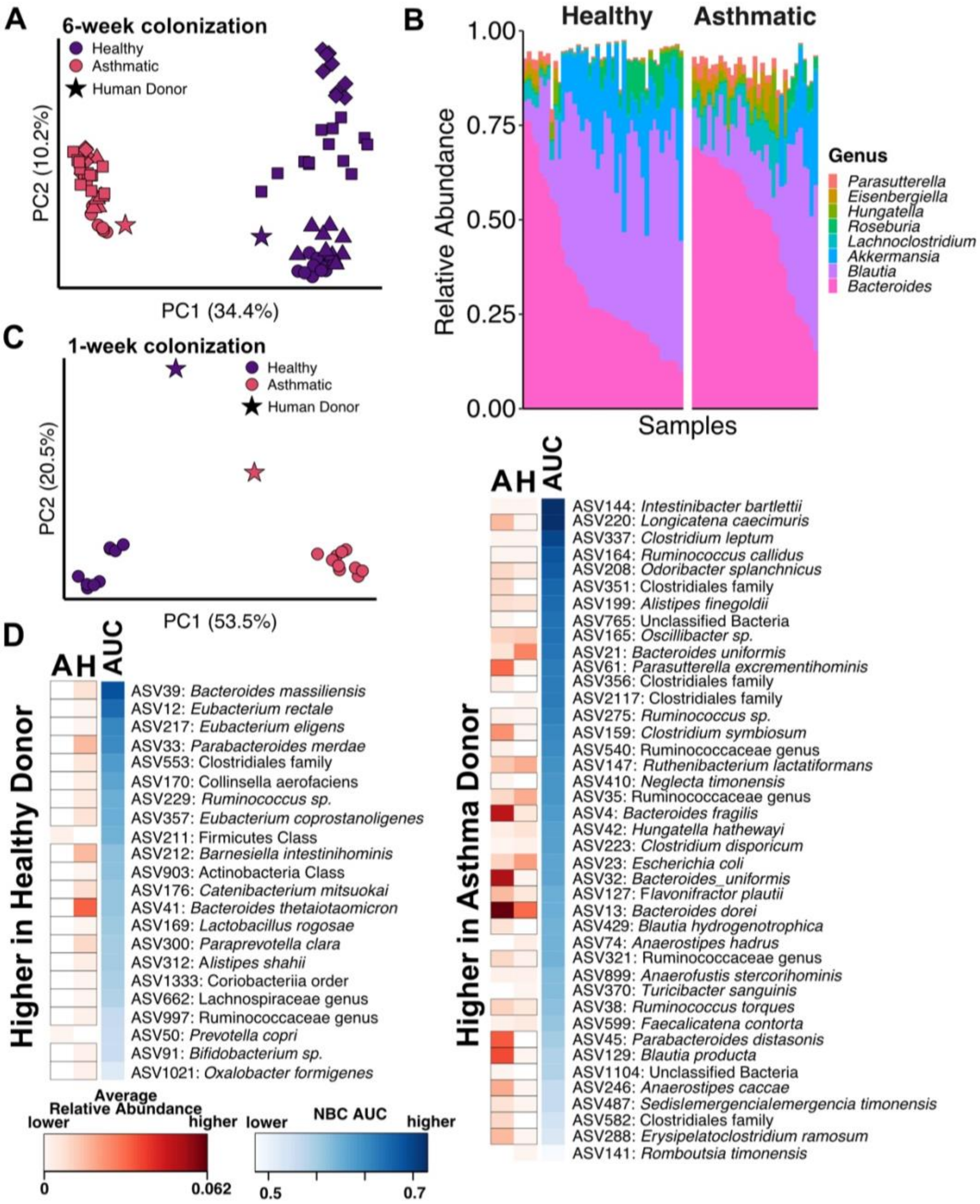


**Supplemental Figure 8: 16S rRNA metagenomic characterization of stool from humanized gnotobiotic mice A)** PCoA of the fecal 16S Unifrac distances of humanized gnotobiotic mice colonized for six weeks (n=5-10/group). Shapes correspond to experiments. Star corresponds to human donor sample. **B)** Relative abundance bar plot of all HO and AO mice colonized for six weeks as part of this study (n=5-10/group, 4 experiments). **C)** PCoA of the fecal 16S Unifrac distances of humanized gnotobiotic mice colonized for one week (n=9-10) only along with donor fecal samples (stars). **D)** Heatmap of taxa concordant by NBC and occurring in HO and AO mice colonized with human donors. Statistically significant differences noted by black rectangle outlines (Wilcoxon, two-tailed).

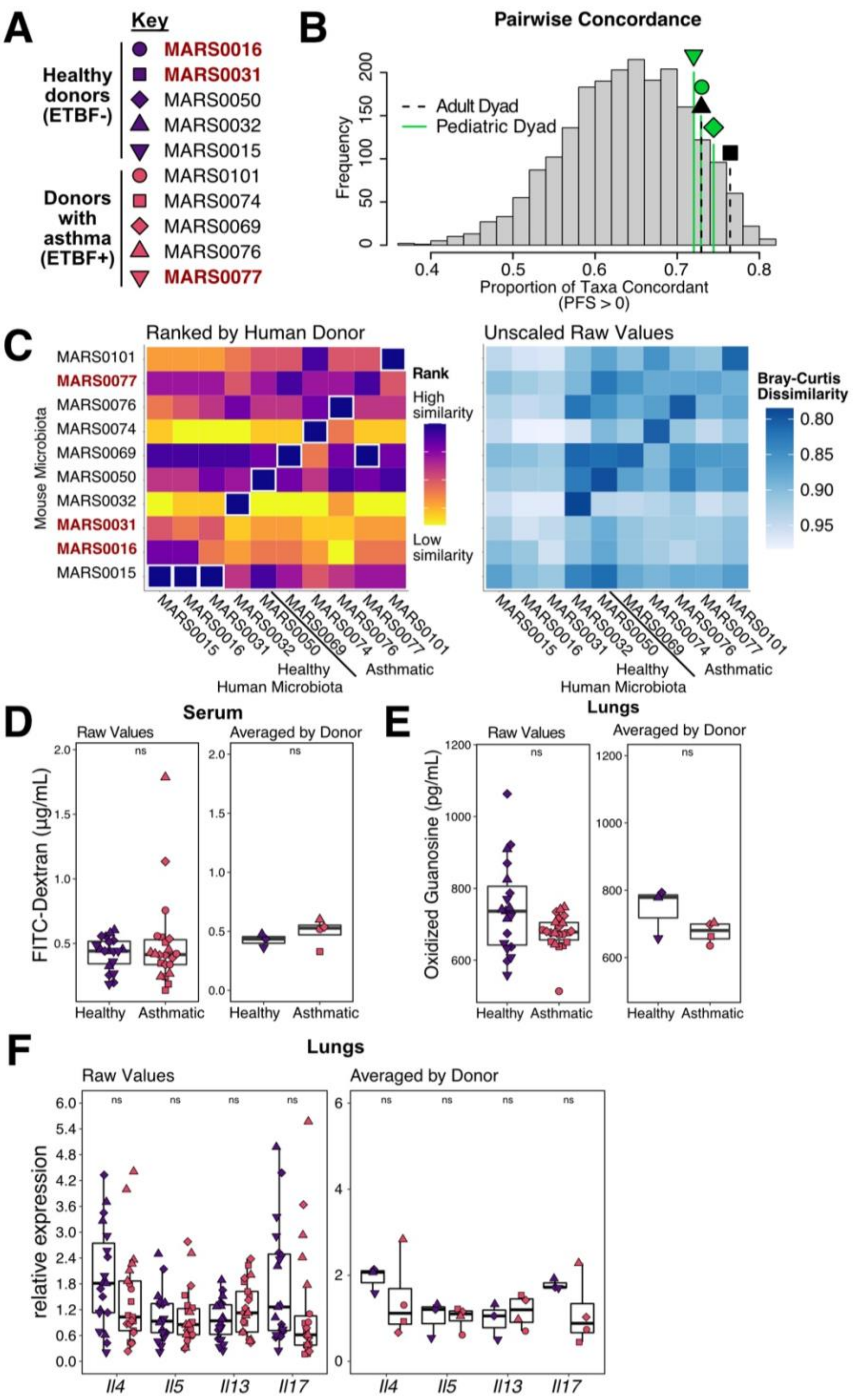


**Supplemental Figure 9: Humanization with additional ETBF+ microbiota does not always increase intestinal permeability or markers of AAI compared to humanization with healthy controls**

**A)** Legend key of MARS donors used for each humanization. Shapes correspond to matched samples by age group (pediatric vs. adult) and microbiome composition. Red names were excluded from analysis. **B)** Histogram of the proportion of pairwise concordant taxa across all possible healthy-asthma donor dyads (also shown in Figure 3C). Vertical dashed line denotes the ETBF+/ETBF- dyads used in this experiment. **C)** Heatmap of Bray Curtis dissimilarity between stool from human donors and recipient mice at the time of sacrifice. The heatmap is presented as ranked across human microbiomes to highlight mouse samples most reflective of their donor (left, top rank outlined in white) and as raw dissimilarity values (right). Mouse recipients in red and bolded were not the most similar to their donor and were excluded from further analysis. **D)** Intestinal permeability of ovalbumin sensitized and challenged (OSC) mice colonized with healthy or asthmatic microbiota following a 2-week colonization (total Healthy n=19, total Asthmatic n = 22). **E)** Oxidized guanosine in lungs of humanized OSC mice F) Lung tissue expression of genes encoding IL-4, -5, -13, and -17A measured by RT-qPCR relative to healthy donors (total Healthy n = 18-19, total Asthmatic n = 21-22)

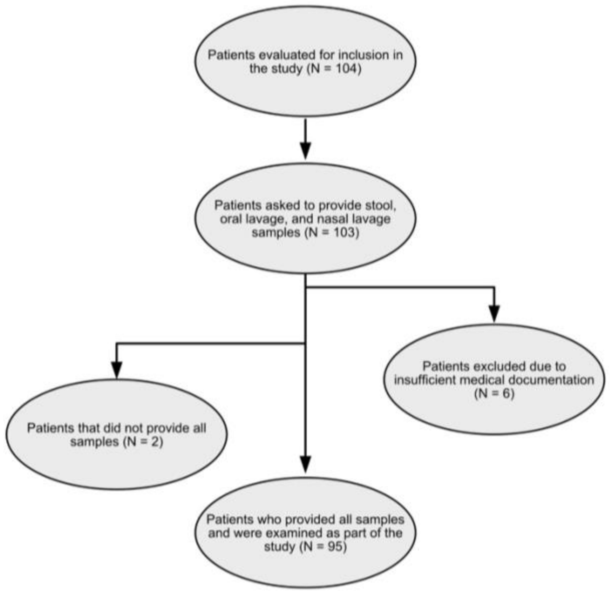


**Supplemental Figure 10: Breakdown of MARS participants inclusion**

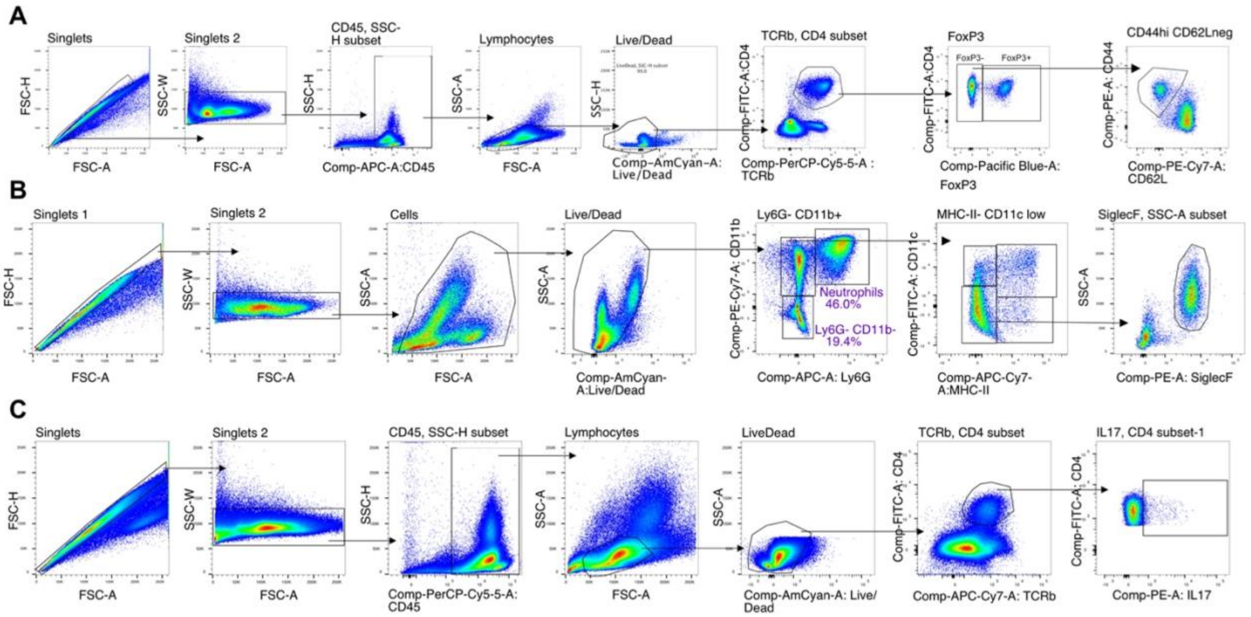


**Supplemental Figure 11: Flow cytometry gating strategies A)** effector T-cells, **B)** neutrophils and eosinophils, and **C)** IL-17A producing effector T-cells.

### Supplemental Tables

**Supplemental Table 1** - Demographics of Participants in Microbiome and Asthma Research Study

**Supplemental Table 2** - Results of PERMANOVA (adonis2) analysis on Bray-Curtis distances of the V4 16S profile of stool from MARS participants

**Supplemental Table 3**: Differentially abundant amplicon sequence variants (ASVs) identified by DESEQ2 along with Naïve Bayes' Classifier (NBC) and Random Forest (RF) Importance

**Supplemental Table 4**: FGSEA analysis of the lungs of mice colonized with either an asthmatic donor or healthy microbiota and sensitized to and challenged with ovalbumin

**Supplemental Table 5**: Summary of all genes in RNA Sequencing of lungs tested using DESEQ2 in one of four comparisons

**Supplemental Table 6**: Amplicon sequence variants (ASVs) identified from the MARS study and their classification using RDP Classifier

**Supplemental Table 7**: Importance and rank of amplicon sequence variants (ASV) included in Naïve Bayes' Classifier (NBC) and Random Forest Model (RF)

**Supplemental Table 8** - Primer Table
